# Dual targeting of GPX4 and TXNRD1 triggers eradication of AML cells through induction of apoptosis and ferroptosis

**DOI:** 10.1101/2024.04.03.584800

**Authors:** Cécile Favreau, Coline Savy, Maxence Bourgoin, Thomas Botton, Sarah Bailly, Florence Granger, Catherine Birck, Marwa Zerhouni, Emeline Kerreneur, Alban Vigroux, Jade Dussart Gautheret, Marie-Laure Arcangeli, Arnaud Porterszman, Thomas Cluzeau, Stephane Rocchi, Arnaud Jacquel, Rachid Benhida, Patrick Auberger, Anthony R Martin, Guillaume Robert

## Abstract

MyeloDysplastic Syndromes (MDS) are a group of heterogeneous hematological disorders characterized by bone marrow failure and abnormal hematopoietic cell expansion, often progressing to acute myeloid leukemia (AML). Current treatments for AML and high-risk MDS have limited efficacy, requiring the exploration of new therapeutic approaches. Recent research highlighted the potential of inducing cell death through ferroptosis, either independently or alongside traditional chemotherapy, as promising approaches for treating MDS/AML cells. We described here two novel compounds, HA344 and #231, capable of targeting both ferroptosis and apoptosis, leading to the effective eradication of cell lines and primary blasts from MDS/AML patients, while sparing normal hematopoietic cells. RNASeq analysis identified oxidation reduction and apoptotic processes as highly significant induced pathways in two different AML cell lines. Using click-chemistry approaches coupled to mass spectrometry, we identified glutathione peroxidase 4 (GPX4) and thioredoxin reductase 1 (TXNRD1) as the main targets of HA344 and #231 in a large panel of AML cells. Accordingly, both compounds inhibited GPX4 and TXNRD1 activity in the micromolar range and triggered GPX4 degradation. Moreover, using recombinant GPX4 carrying or not a selenium (GPX4-Se and GPX4-S), we confirmed by mass spectrometry that HA344 and #231 bind more efficiently GPX4-Se than GPX4-S. In conclusion, these compounds might represent a new pharmacological approach in the treatment of MDS and AML, offering a potential avenue for future therapies.

## INTRODUCTION

MyeloDysplastic Syndromes (MDS) consist of a heterogeneous spectrum of myeloid clonal hematological disorders that appear frequently in elderly patients with a median age at diagnosis of about 70 ^1^. This disease which represents about 90.000 new cases of malignant neoplasms every year worldwide incorporates 2 major clinical features: a differentiation defect of hematopoietic cells that results in bone marrow failure associated with peripheral blood cytopenias, and expansion of at least one abnormal hematopoietic clone which undergoes, in about 30% of MDS cases, evolution to acute myeloid leukemia (AML) ^2,3^. Recent high throughput sequencing-based analyses allowed a better characterization of MDS, that was referred as a pre-leukemic disease in which most of the mutations recurrently identified were gained in pre-leukemic genes such as TET2, SF3B1, SRSF2, ASXL1, RUNX1, DNMT3A, and EZH2 to promote clonal expansion of pre-leukemic MDS cells ^4,5^. Eventually, another subset of mutations is acquired thereafter to promote progression to AML ^6^. Therefore, both MDS and AML diseases display a wide heterogeneity of morphologic, cytogenetic, and clinical features which limit the ability of clinicians to make a clear risk assessment. Consequently, treatments for AML and higher-risk MDS in patients younger than 65 have remained largely unchanged for the past four decades. This includes a dose-intensive chemotherapy based on combination of anthracyclines with cytarabine (Ara-C), and for eligible patients, bone marrow transplantation ^7^. Given the high toxicity of AML-like chemotherapy in older patients with higher-risk MDS, Azacitidine and Azacitidine + Venetoclax has now replaced the anthracycline/Ara-C combination ^8^. These agents can achieve a sustained remission in about 70% of cases and even a cure in some patients; however, almost all patients relapse with a progressively more chemo-resistant disease. The mechanisms underlying this resistance process remain a conundrum and are currently subjected to intense research in the field of hematology, not only for fundamental biological insights but also to find an appropriate therapeutic strategy to circumvent durably the relapsed disease. In this line, identifying new molecules able to eradicate MDS and AML cells is a major challenge. To be efficient as potential future therapies these molecules should 1) affect preferentially MDS and AML cells, sparing normal hematopoietic cells and 2) be able to inhibit cell growth or/and induce cell death.

Different forms of cell death have been characterized so far, including apoptosis, necroptosis, pyroptosis, autophagic cell death and more recently ferroptosis ^9–16^. Briefly, apoptosis is a selective mode of programmed cell death executed by a highly conserved family of cysteine proteases, called caspases. Apoptosis can be induced by specific cell death receptors, such as Fas through the extrinsic pathway or by the intrinsic pathway that is regulated by the members of the Bcl2 family ^17^. Both pathways culminate in caspase activation and the cleavage of cellular substrates that elicit the apoptotic response. Reinducing apoptosis in MDS/AML cells using BH3 peptidomimetics, such as Venetoclax, represented a major advance in the management of elderly patients suffering these diseases^18^.

In addition, ferroptosis describes a novel mode of cell death induced by Erastin an inhibitor of the Xc transport system or RSL3 an inhibitor of GPX4 ^19^. The hallmarks of ferroptosis include accumulation of membrane lipid hydroxy peroxides, increased intracellular reactive oxygen species, oxidative stress, and mitochondrial shrinkage. Thus ferroptosis can be defined as a non-apoptotic mode of cell death, driven by iron accumulation and membrane lipid peroxidation when GPX4 activity is inhibited ^20–22^.

Many cancer cell lines displayed dysregulation of one or several cell death mechanisms that contribute to their resistance to different types of drugs. Reactivation of selective modes of cell death in these tumor cell could thus represent a promising therapeutic option to treat different types of cancers. In line with this strategy, we described here two hit compounds HA344 and #231, identified and validated based on a structure-activity analysis of over 200 synthesized molecules. Both compounds were shown to efficiently kill a panel of AML cell lines and bone marrow cells isolated from MDS and AML patients, while sparing normal hematopoietic cells. Both compounds triggered caspase activation and lipid hydroxy peroxidation, culminating in apoptosis and ferroptosis. Consequently, concomitant blockade of apoptosis and ferroptosis abrogated the effects of HA344 and #231. In addition, RNAseq analysis highlighted two main pathways contributing to the effect of both compounds, namely apoptosis and oxidative stress. Accordingly, HA344 and #231 activated NRF2 and triggered a protective response through the induction of genes containing antioxidant response elements (AREs). *In cellulo* click-chemistry experiments coupled to mass spectrometry in different AML cell lines, allowed the identification of 22 common targets among which two selenoproteins TRNXD1 and GPX4 involved in ferroptosis regulation. HA344 and #231 were identified as direct inhibitors of GPX4, and invalidation of GPX4 in AML cell lines phenocopied the effects of the drug.

In conclusion, we described here a new pharmacological approach with two related small covalent inhibitors to eradicate MDS and AML cells through the concomitant activation of apoptosis and ferroptosis. Both compounds exhibit an original mechanism of action by directly targeting and degrading key enzymes involved in the protection of leukemic cells from apoptosis and ferroptosis.

## RESULTS

### HA344 and #231 induce cell death of AML cell lines and efficiently kills CD34^+^ cells from AML patients

We previously described that HA344 (**Fig. S1a**) was highly effective on human melanoma cell lines ^23^. To evaluate the efficacy of HA344 and #231, a nucleoside analog of HA344 (**Fig. S1b**) we performed NCI-60 screening of the 60 most common cell lines representative of cancers from all tissues (**Fig. S1c**). Compound #231 proved particularly effective in inhibiting growth of different leukemic cell lines in the 10^-7^-10^-5^M range, while most of solid cancer cell lines were 10-fold less sensitive to the drug. To further evaluate the efficiency of HA344 and #231 on leukemic cells, we performed dose response curves for these two compounds on 5 AML cell lines representative of the pathology and harboring different types of chromosomal translocation and/or mutations: HL-60 (CDKN2A, NRAS, TP53), OCI-AML3 (NPM1, DNMT3a), MV4-11 (FLT3-ITD, MLL-AF4), NB4 (PML-RARa), and MOLM14 (FLT3-ITD, MLL-AF9). Using DAPI staining **(Fig. 1a)** and XTT assay **(Fig. 1b)** we observed that HA344 and #231 compounds killed AML cells very efficiently. Nevertheless, both compounds were slightly more efficient on MOLM-14 and MV4-11 AML cells, that harbor FLT3-ITD mutations. Importantly, peripheral blood mononuclear cells (PBMC) from healthy donors treated with increasing doses of HA344 and #231 were less sensitive to cell death induction after a 48h treatment **(Fig. 1c)**. The LD50 values for HA344 and #231 in different AML cell line ranged from 0.4 to 1.3 μM but were 10 to 40 times higher in PBMC from healthy patients **(Fig. 1d)**.

**Figure 1.**
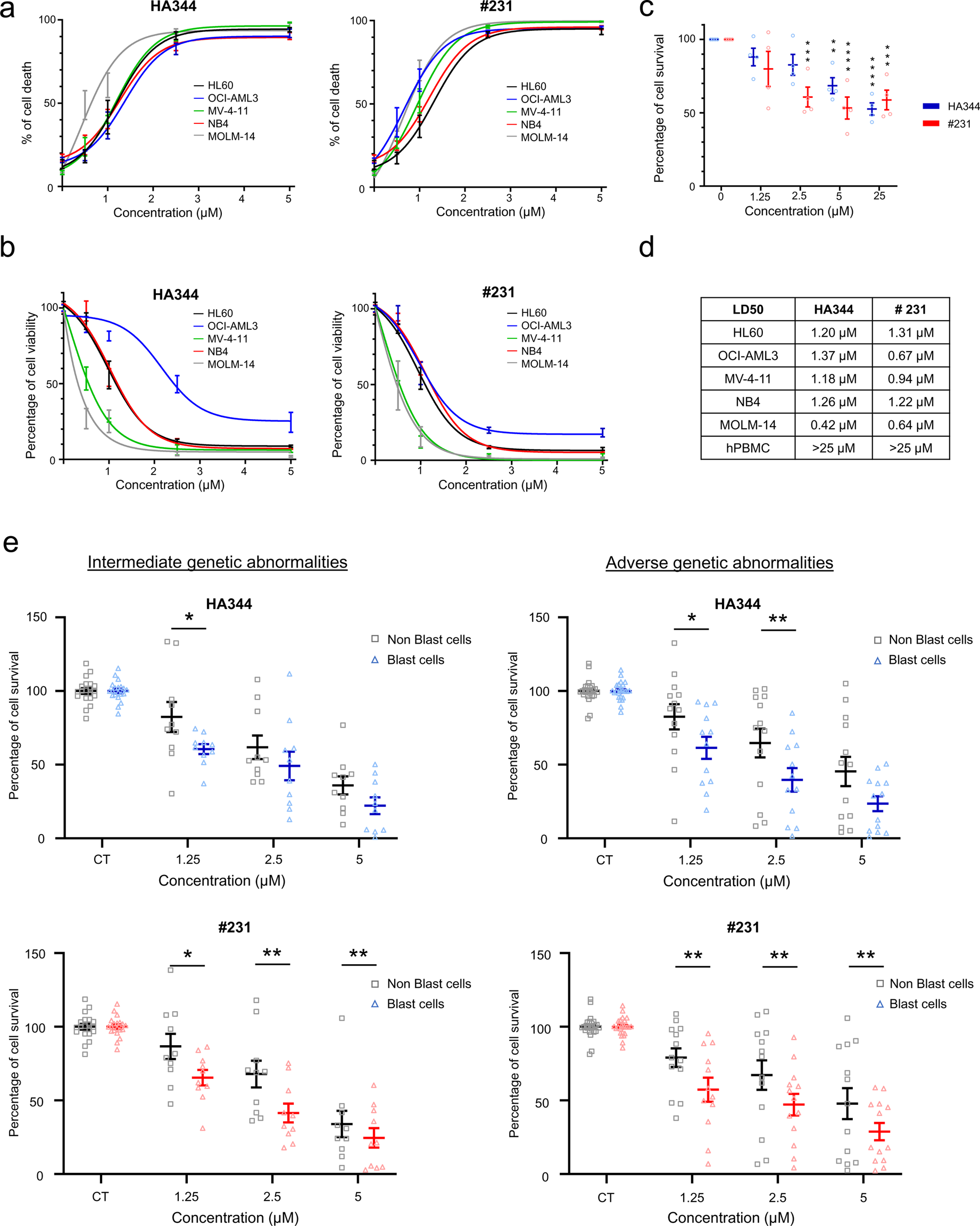
HA344 and #231 promote cell death of leukemia cells. HL60, OCI-AML3, MV4-11, NB4, and MOLM-14 cell lines were exposed to increasing concentrations of HA344 and #231, ranging from 0.5 µM to 5 µM. After a 48-hour incubation period, cell death was evaluated using **(a)** Dapi incorporation and **(b)** cell viability assays using the XTT assay and fluorescence readings. Healthy donor-derived peripheral blood mononuclear cells (hPBMC) were treated with HA344 and #231 for 48 hours, and **(c)** cell survival was assessed using flow cytometry by analyzing Dapi exclusion. **(d)** Lethal half-doses 50 (LD50) for each of the cell lines and for hPBMC. Bone marrow (BM) cells obtained from patients were stratified into two groups based on their genetic status namely “Intermediate genetic abnormalities” and “Adverse genetic abnormalities. BM cells were exposed to HA344 and #231 at concentrations of 1.25 µM, 2.5 µM, and 5 µM. Following a 48-hour treatment period, blast cells and non blast cells were selectively identified, and **(e)** cell survival was evaluated using flow cytometry by the exclusion of Dapi incorporation.

To further investigate the anti-leukemic effect of our molecules, 27 individual medullary biopsies patients diagnosed with MDS/AML were treated with increasing concentrations of HA344 or #231 for 48h (**Table S1**). Among them, 12 patients had intermediary gene abnormalities and 15 had adverse genetic abnormalities. Cell death induced by HA344 and #231 was evaluated on hematopoietic cells (CD45+), as well as blast cells (CD45+/CD34+) using Dapi staining, as illustrated in **Supplemental Figure 1d.** HA344 and #231 triggered a dose-dependent increase in cell death in bone marrow samples from AML patients. Of greater interest was the fact that blast cells displayed a statistically significant increase in their sensitivity to both molecules when compared to non-blast cells. Importantly, this effect was observable in patients with both intermediate and adverse genetic abnormalities **(Fig. 1e**).

### HA344 and #231 induce transcriptional modifications in the apoptotic and ROS-dependent pathways, contributing to the elimination of malignant myeloid cells

To go further into the mechanism of action of our compounds, we conducted an analysis of mRNA expression using NGS RNA-seq on both HL-60 and OCI-AML3 cells following a 10h treatment with either HA344 or #231. We first illustrated the sample distances for both control and treatment conditions relative to each cell line and compound (**Fig. S2a-d-left panel**). These analyses revealed the remarkable homogeneity of our samples, while demonstrating that HA344 and #231 leads to a significant regulation of mRNA expression as illustrated by the volcano plots presented in (**Fig. S2a-d-right panel**). Notably, upregulated genes exhibited a higher level of regulation compared to downregulated ones. In HL-60 cells, HA344 treatment resulted in a seven-fold increase in upregulated genes compared to downregulated ones, while in OCI-AML3 cells, it induced a two-fold increase as compared to downregulated ones. Likewise, compound #231 induced a threefold increase in upregulated genes compared to downregulated ones in HL-60 cells and a twofold increase in OCI-AML3 cells. Of note, the extent of gene regulation was substantially greater in HL-60 cells, where HA344 influenced the regulation of 7396 genes and #231 affected 17589 genes, contrasting with OCI-AML3, where HA344 affected 3121 genes and #231 affected 3611 genes (**Fig. S2a-d-right panel**). Using the Gene Ontology classification system, we identified apoptotic death process and response to oxidative stress as the most prominently regulated pathways among the 17 most regulated pathways on a total panel of 9403 (**Fig. 2a)**. Further analysis of the GSEA results established that the NRF2 pathway was also highly enriched in HL60 and OCI-AML3 treated cell lines. The normalized enrichment scores (NES) for HL60 cells treated with HA344 and #231 were 1.55 and 1.29, while in OCI-AML3 cells, the scores were 1.51 and 1.71, respectively, demonstrating the significant induction of NRF2-dependent genes occurs upon treatment with HA344 and #231(**Fig. 2b**). All together these data are compatible with a major effect of our compounds on the regulation of the apoptotic death process and the response to oxidative stress.

**Figure 2.**
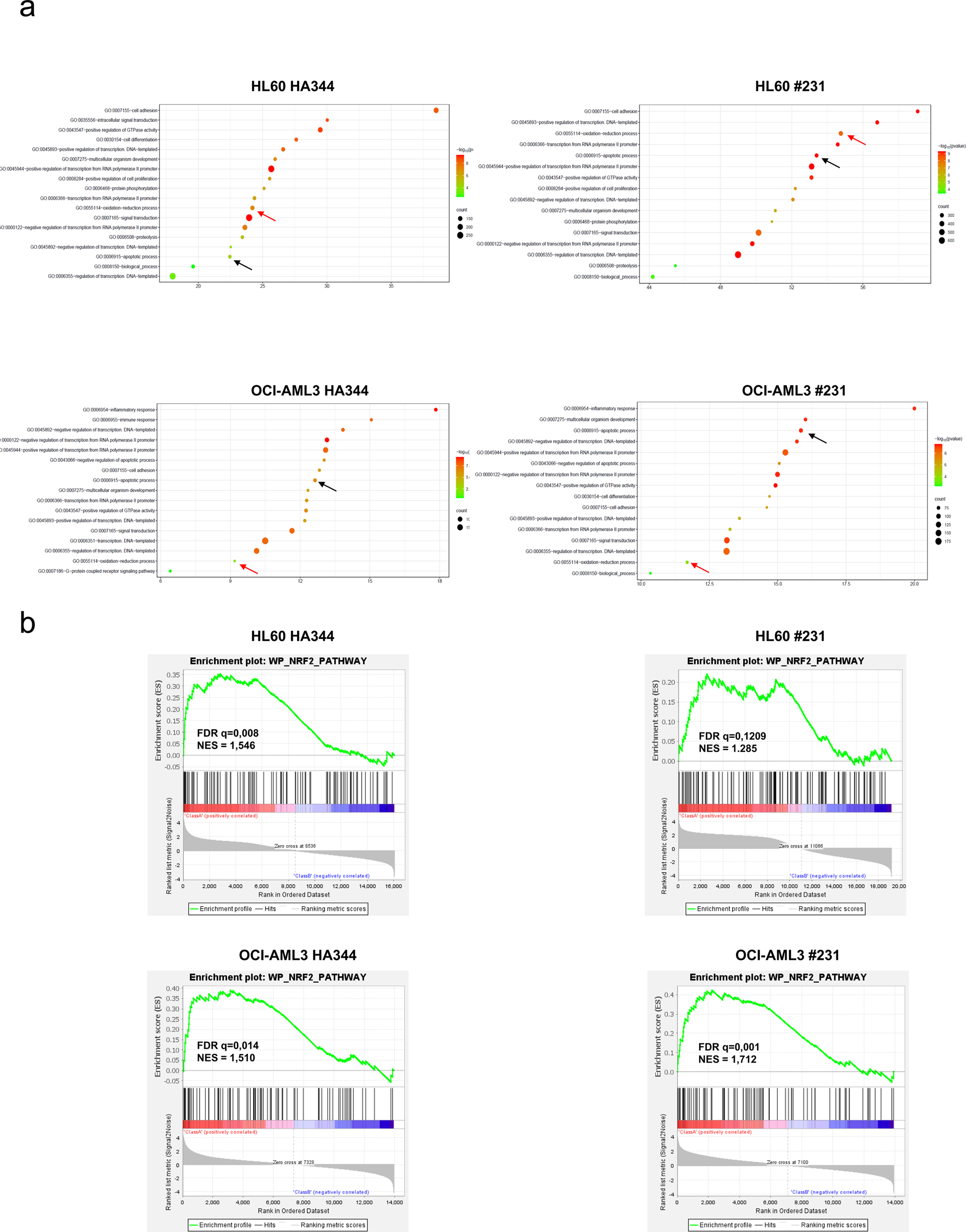
HA344 et #231 induce both apoptotic and ROS-dependent cell death. (a) HL60 and OCI-AML3 were treated 6h with LD50. RNA-seq data collected from HL-60 and OCI-AML3 cells after exposure to HA344 and #231 at their respective DL50 concentrations for 6 hours were examined using the Gene Ontology (GO) approach. The results show the top 17 Gene Ontology (GO) categories ranked according to the proportion of significant genes. The degree of significance is indicated by color, and the size of the circle reflects the number of significant genes in each category. **(b)** Gene Set Enrichment Analysis (GSEA) was conducted in two groups: CT vs. HA344 (left panels) and CT vs. #231 (right panels). The GSEA algorithm computes an enrichment score (ES), which signifies the level of overrepresentation at the upper or lower portion of the ranked list of genes within a specific gene set, as compared to all genes in the RNA-seq dataset. A positive ES indicates an enrichment of the gene set at the top of the ranked list. The analysis reveals that the NRF2 pathway is enriched in both the HA344 and #231 treated groups.

### Induction of cell death by HA344 and #231 is inhibited by apoptosis and ferroptosis inhibitors

In view of the results obtained in the RNA-seq analysis, HL-60 and OCI-AML3 cell lines were subjected to pretreatment with the pan-caspase inhibitor Q-VD-OPh (QVD), followed by treatment with HA344 or #231 at their LD50 concentrations for 24 h or 48 h **(Fig. 3a)**. HA344 and #231-induced cell death was significantly reduced in the presence of QVD, both in HL-60 and OCI-AML3 cells at 48h **(Fig. 3a)**. To further support the involvement of apoptosis in the cell death mechanism induced by our compounds, we carried out caspase activity assays. Both HA344 and #231 induced a robust activation of initiator caspases 3 and 7 **(Fig. 3b, left panel)** as well as effector caspase 9 **(Fig. 3b, right panel)** after 24 h of treatment. These data confirm the involvement of caspases in the cell death process induced by the two compounds. To strengthen our findings, HL-60 cells were exposed to HA344 or #231 at concentrations equivalent to 1 or 2 times the LD50 for 24 h **(Fig. S3a)**. Western blot analysis confirmed the cleavage of procaspase 3 and PARP. Indeed, we observed that at the higher concentration of both compounds (2x LD50) the zymogen of caspase 3 completely disappeared. These findings provide robust evidence supporting the implication of caspase and apoptosis in HA344 and #231 effects.

**Figure 3.**
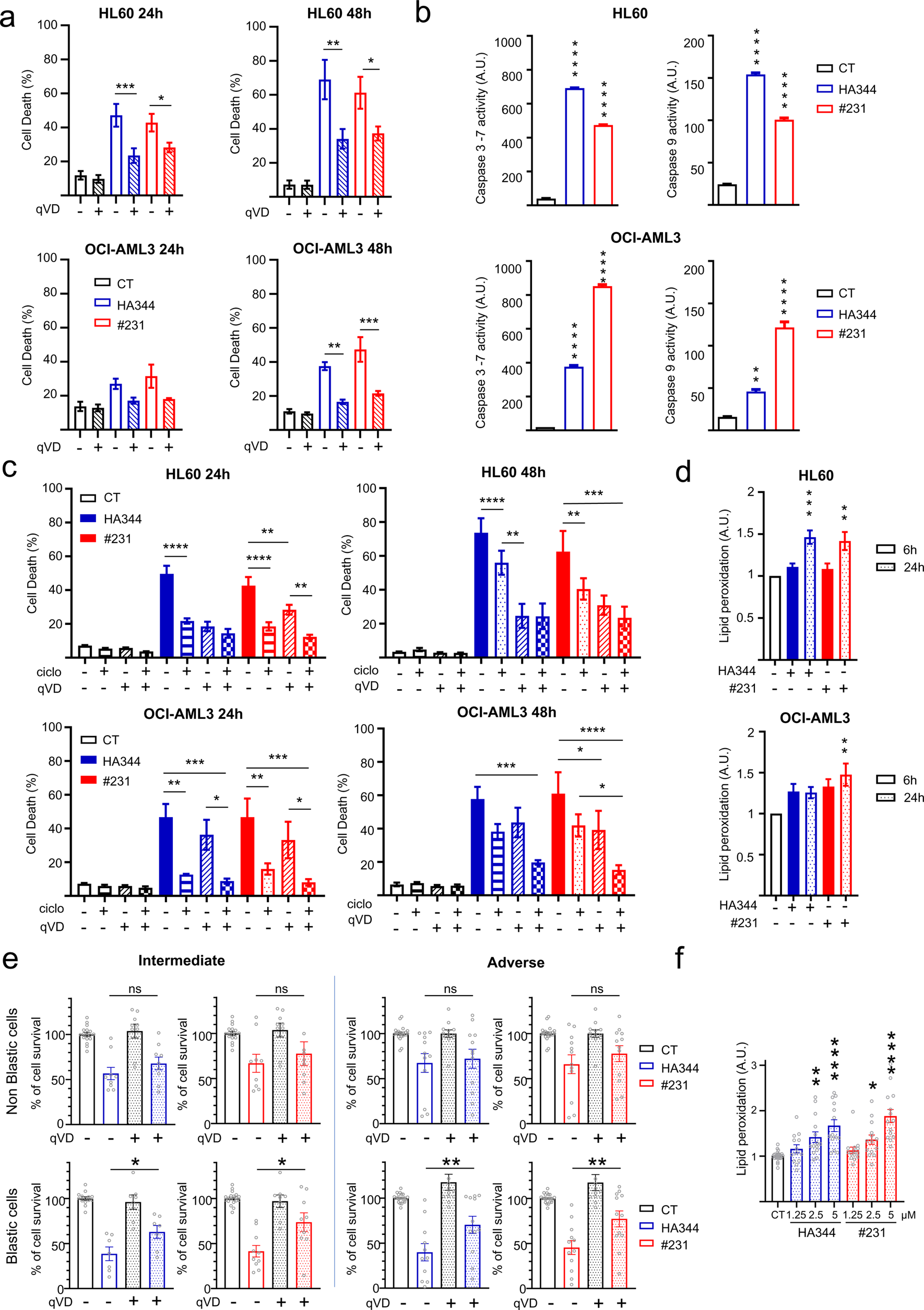
HA344 and #231 induce both apoptosis and ferroptosis in leukemia cells. (a) HL60 and OCI-AML3 cell lines were pretreated with 20 µM qVD, before the addition of HA344 or #231 at their respective LD50 concentrations. Cell death was assessed at 24 and 48 hours by measuring DAPI incorporation. **(b)** After 24 hours of treatment with HA344 and #231 at their LD50 levels, Caspase 3-7 and 9 activities were measured in the HL60 and OCI-AML3 cell lines. **(c)** HL60 and OCI-AML3 cell lines were pretreated for 1h with either 20 µM qVD, 1 µM ciclopirox, or a combination of both, followed by incubation with HA344 and #231 at their LD50 concentrations. Cell death was quantified by measuring DAPI incorporation at 24 and 48 hours. **(d)** Lipid peroxidation was evaluated in HL60 and OCI-AML3 cells treated with HA344 and #231 at their LD50 concentrations. Assessment was conducted at 6 and 24 hours using a specific probe (Lipid peroxidation assay kit). **(e)** Patient BM samples were pretreated for 1h with 20 µM qVD and then incubated with 2.5 µM of HA344 or #231. After 48 hours, blast cells were distinguished from non-blast cells, and cell death was assessed by DAPI incorporation. **(f)** Patient BM samples were treated with 1.25, 2.5, and 5 µM of HA344 or #231. After 24 hours, the quantification of peroxidized lipids was performed using a Lipid Peroxidation Assay, followed by flow cytometry analysis.

In line with the results of our RNA-seq analysis, we next used a large-scale pharmacological approach to inhibit various cell death processes or ROS generation. For the sake of simplicity, we only depicted the results with the two most active inhibitors i.e QVD and ciclopirox, an iron chelator. Both inhibitors were found to statistically inhibit the antileukemic effects of HA344 and #231 at 24 and 48h (**Fig. 3c)**. Combination of QVD and ciclopirox abrogated the effect of HA344 and #231 on cell death **(Fig. 3c),** suggesting that apoptosis and ferroptosis both contributed to the effects of our compounds. We finally confirmed the inhibition of HA344 and #231-induced cell death using deferoxamine another iron chelator and known inhibitor of ferroptosis **(Fig. S3b)**. Ferroptosis is a recently recognized form of regulated cell death triggered by lipid peroxidation of membrane proteins and characterized by iron dependency. The involvement of ferroptosis in addition to apoptosis in HL-60 and OCI-AML3 cells was confirmed by the increase in lipid peroxidation at 6 or 24h of HA344 and #231 treatment (**Fig. 3d**). Moreover, increasing the concentrations of both compounds clearly triggers a dose-dependent augmentation of lipid peroxidation **(Fig. S3c)**. We next labelled cells with a fluorescent probe undergoing a color shift from red to green, to perfom a ratiometric measurement of lipid peroxidation. Following a 3h treatment with HA344, notable changes including condensation were observed in the nucleus and cytoplasm of the HL60 cell line. Moreover, there was a noticeable enhancement in the intensity of green fluorescence, which reflects an increase in lipid peroxidation **(Fig. S3d)**. Pictures in **Fig. S3d** also showed the appearance of circular membrane protrusions, known as macro-blebbing, a characteristic feature of ferroptosis ^24^.

Having established the role of apoptosis in HA344 and #231 effects in myeloid leukemia cell lines, we next pretreated BM-derived blastic and non-blastic cells from AML patients to QVD, followed by a 24h treatment with HA344 or #231. QVD was shown to mitigate HA344 and #231 mediated cell death, mainly within blast cells population **(Fig. 3e)**, thereby validating the importance of apoptosis induction by our compounds in patient blasts. Additionally, we evaluated lipid peroxidation levels in patient samples following a 24h treatment with increasing concentrations of HA344 or #231 **(Fig. 3f)**. Like with AML cell lines, we detected a significant increase in lipid peroxidation in cells from AML patients, thus confirming the involvement of ferroptosis in the mechanism of action of our compounds in patient’s cells.

### HA344 and #231 trigger ROS production, NRF2 relocation and overexpression of genes containing ARE elements

Considering the effect of our compounds on lipid peroxidation we analyzed their effect on ROS production using the DCFDA probe. Treatment of AML cell lines with HA344 and #231 triggered a comparable induction of ROS at 30 min (**Fig. 4a and 4b**). However, ROS induction was less sustained in OCI-AML3 cells compared to HL-60 cells. These findings are in line with the lower sensitivity of OCI-AML3 cells to both drugs **(Fig. 1b)**. As GSEA analysis previously established a significant induction of the NRF2 signaling pathway under HA344 and #231 treatment **(Fig. 2b)**, AML cell lines were treated for 10h with HA344 and #231 and cytoplasmic and nuclear extracts were prepared and subjected to Western Blot analysis. Both compounds were found to trigger the nuclear accumulation of the transcription factor NRF2 **(Fig. 4c)**. As accumulation of NRF2 in the nucleus is associated with increased gene expression related to the antioxidant response, we performed RT-qPCR analysis of well characterized antioxidant genes NRF2, NADPH quinone dehydrogenase 1 (NQO1), heme oxygenase 1 (HMOX1), thioredoxin (TXN), Thioredoxin reductase 1 (TXNRD1), TXNRD2, glutamate-cysteine ligase catalytic subunit (GCLG) and glutamate-cysteine ligase modifier subunit (GCLM) in AML cell lines treated with HA344 and #231 **(Fig. 4d-e and Fig. S4b-c)**. In HL60 cells, HA344 and #231 drastically increased the expression of HMOX1 mRNA (400 to 600-fold) at 6h while expression of NRF2, NQO1 TXNRD1 and GCLM increased more moderately (2 to 4 times) **(Fig. 4d-e)**. We observed highly similar results in OCI-AML3 AML cells **(Fig. S4b-c)**. All together, these observations clearly indicate that H344 and #231 trigger NRF2 nuclear relocation leading to increased expression of ARE responsive genes. A list of additional genes linked to the NRF2 signaling pathway is provided in **(Fig. S5)**.

**Figure 4.**
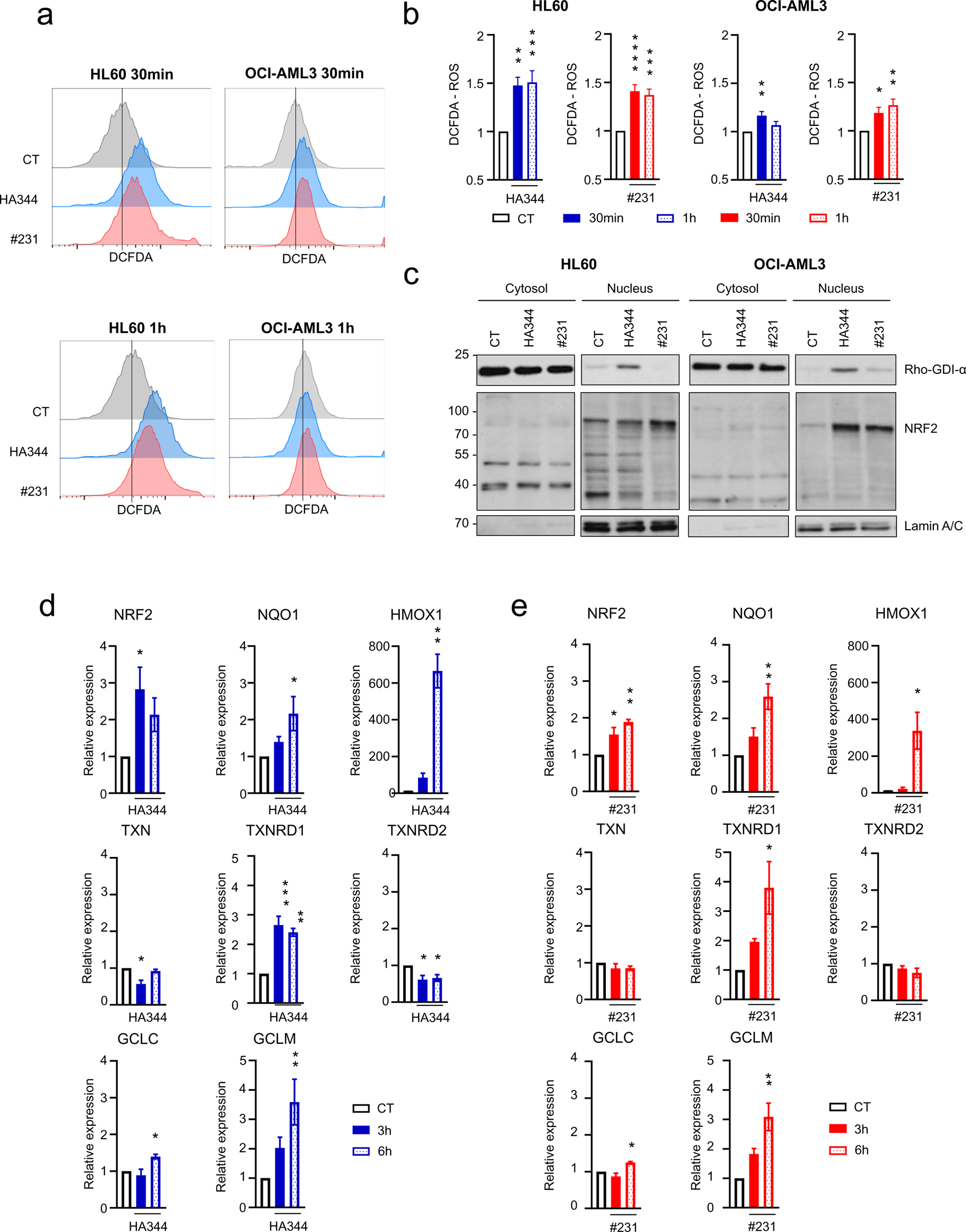
HA344 and #231 compounds elevate reactive oxygen species (ROS) levels in leukemia cell lines. (a) HL60 and OCI-AML3 cell lines were treated 30 min or 1h with HA344 and #231 at the LD50, then ROS levels were assessed by flow cytometry analysis using the DCFDA probe. **(b)** The histograms depict the mean fluorescence intensity of the quantified ROS **(c)** HL60 and OCI-AML3 cell lines were subjected to a 10-hour treatment with HA344 and #231 at LD50. NRF2 translocation from the cytosol to the nucleus was monitored through subcellular fractionation and analyzed by western blotting. **(d-e)** Analysis of the mRNA expression levels of NRF2-dependent genes using RT-QPCR in HL60 cells after treatment with HA344 or #231 at LD50 for either 3 or 6 hours.

### Identification of GPX4 as a substrate for HA344 and #231 compounds in AML cell lines

Identification of cellular targets remains a challenge for the development of molecules with high anticancer potential. The development of click-chemistry over the last 10 years has greatly improved the discovery of protein targets ^25^. To identify the cellular targets of HA344 and #231 in AML cell lines, we generated the active probes VG41 and VG40, along with their counterparts VG43 and VG60, which do not contain the ethyl propiolate active motif highlighted in red in **Figure 5a**. We first checked that increasing concentrations of VG43 and VG60 failed to induce cell death in three different AML cell lines, namely HL60, OCI-AML3 and MV4-11 after 48h of incubation, conversely to VG41 and VG40 which triggered cell death in all three cell lines **(Fig. S6a)**. We next incubated these three AML cell lines with 2 µM of VG41, VG40 or VG43 and VG60 as negative controls. Following a click reaction with a biotin label, several washing and enrichment steps, eluates were submitted to mono-dimensional gel electrophoresis, staining and finally subjected to mass spectrometry analysis (Please see **Fig. S6b** for the complete protocol).

**Figure 5.**
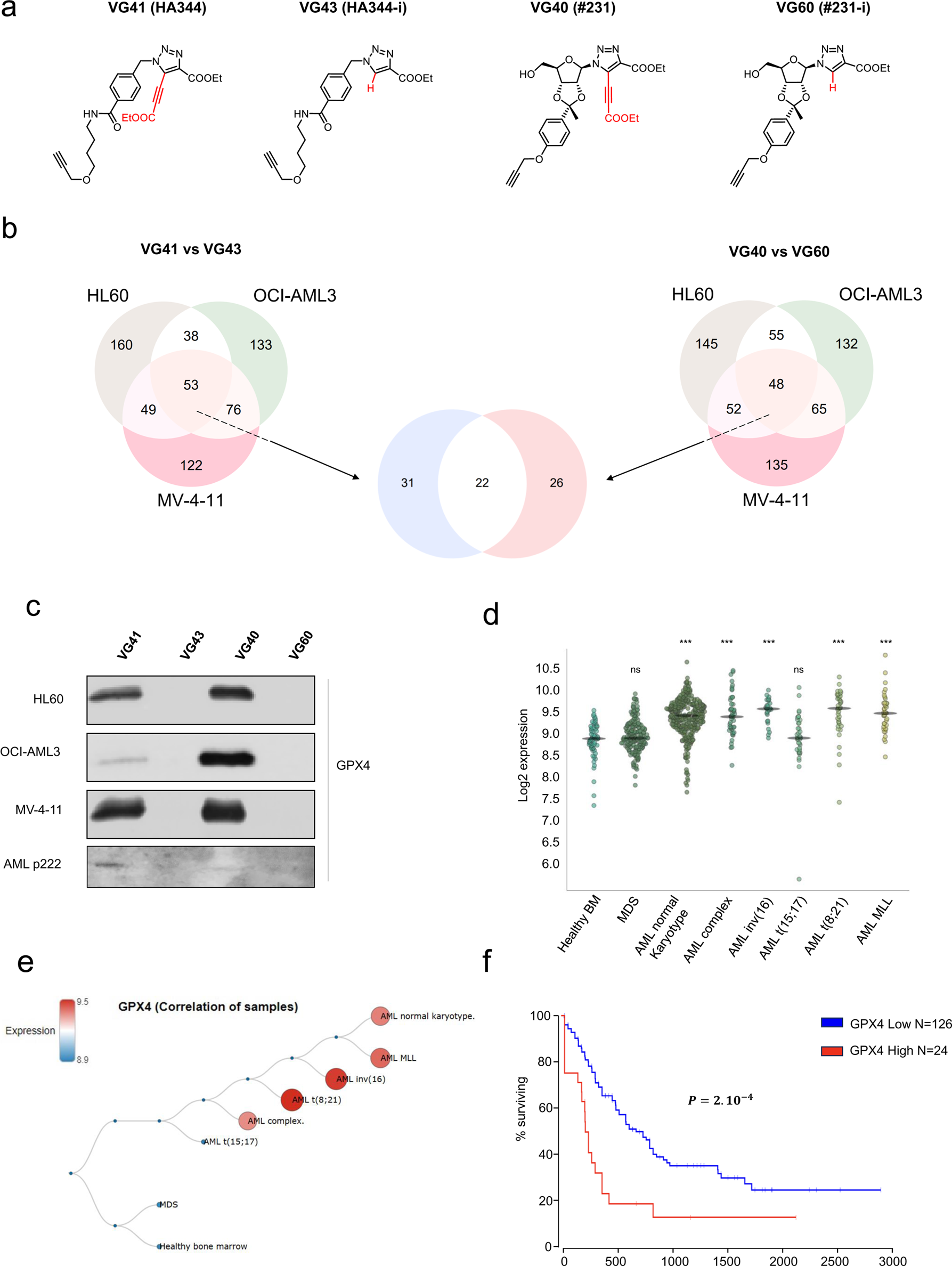
HA344 and #231 mainly target GPX4. (a) Structure of compounds VG41, VG43, VG40 and VG60, representative of the clickable forms of HA344 and #231 with and without the activity-bearing motif. **(b)** The proportional Venn diagrams determine the number of common proteins captured following the application of click chemistry and Mass spectrometry to HL60, OCI-AML3, and MV-4-11 cell lines treated with 2.5 µM of VG41, VG43, VG40, and VG60 over a period of 1 hour. **(c)** HL60, OCI-AML3, MV4-11 cell lines, were subjected to treatment with 2.5 µM of VG41, VG43, VG40, and VG60 for 1 hour, followed by a click chemistry reaction and Western Blot analysis to confirm the binding of HA344 and #231 to GPX4. **(d)** Graph displaying the average GPX4 mRNA expression in various leukemia types, using data sourced from the Microarray Innovations in Leukaemia (MILE) study. **(e)** Hierarchical structure visualizes the relative GPX4 levels in distinct AML subtypes based on their karyotypes, with data also originating from the MILE study. **(f)** The Kaplan-Meier plot is derived from data provided by the Oncolnc website, which conducts analyses on the TCGA cohort. This graph illustrates the comparison of overall survival between patients who demonstrate high GPX4 expression and those with low GPX4 expression levels.

An unbiased analysis of the MS results identified a strong overlap of target proteins between VG41 and VG40 in all cell lines **(Fig. 5b)**. We next examined the common protein targets between VG41 vs VG43 and VG40 vs VG60 **(Fig. 5b)** and identified 53 common targets for VG41 and 49 common targets for VG40 in the three cell lines treated. Among these protein targets,

22 were shared by VG41 and VG40 in the three treated cell lines **(Fig. 5b, lower panel).** The 22 common targets ranked as a function of the enrichment value in the three cell lines, are presented in **Fig. S6c**. For convenience a detailed list for the 22 common targets is presented in **(table S2 and S3)**. Among the 3 AML cell lines tested the most relevant substate for VG41 and VG43 was glutathione peroxidase 4 (GPX4). *The in-cellulo* click-chemistry experiment confirmed that both molecules bind directly to GPX4 in the three AML cell lines **(Fig. 5c)**. Our findings were extended to a patient sample (p222) with AML containing 80% blasts, demonstrating that VG41 and VG43 also trapped GPX4 under identical conditions **(Fig. 5c)**. Data sourced from the Microarray Innovations in Leukemia (MILE) study highlighted an increased in GPX4 mRNA levels across various subtypes of AML, such as those with a normal karyotype, complex karyotype, inv(6), t(18:21), and MLL rearrangement, but not in MDS and AML t(15:17)**(Fig. 5d and e)**. Finally, Kaplan-Meier plots derived from data provided by the oncolnc website (TGCA cohort, NEJM 2012) ^26^ identified high GPX4 mRNA expression as a pejorative prognosis marker in AML patients **(Fig. 5f)**.

### The interaction of HA344 and #231 with GPX4 causes suppression of its activity and triggers its degradation

GPX4 protein levels were evaluated in the 5 AML cell lines already tested for their sensitivity to HA344 and #231. The expression level of GPX4 was roughly similar in HL-60, OCI-AML3, NB4 and MOLM-14 cell lines, while MV4-11 exhibited a significantly higher GPX4 expression level **(Fig. 6a)**. We decided to analyze whether GPX4 knock down might affect cell death and ferroptosis of OCI-AML3 cell line. Transient transfection of GPX4 siRNA for 48h resulted in a drastic inhibition of GPX4 expression in the OCI-AML3 cell line **(Fig. 6b, upper panels)**. Abolition of GPX4 expression triggered a strong increase in lipid peroxidation and induction of cell death, that were abolished or inhibited, respectively in the presence of alpha-tocopherol, a free radical inhibitor known to protect cell membranes from lipid peroxidation **(Fig. 6b, lower left panel)**. Next, we pretreated AML cell lines with alpha-tocopherol before exposing them to HA344 and #231 **(Fig. S7a)**. Alpha-tocopherol significantly reduced cell death induced by our compounds at 24 and 48 h in both HL-60 and OCI-AML3 cells **(Fig. S7a)** to the same extent than GPX4 invalidation **(Fig. 6b, lower right panel)**

**Figure 6.**
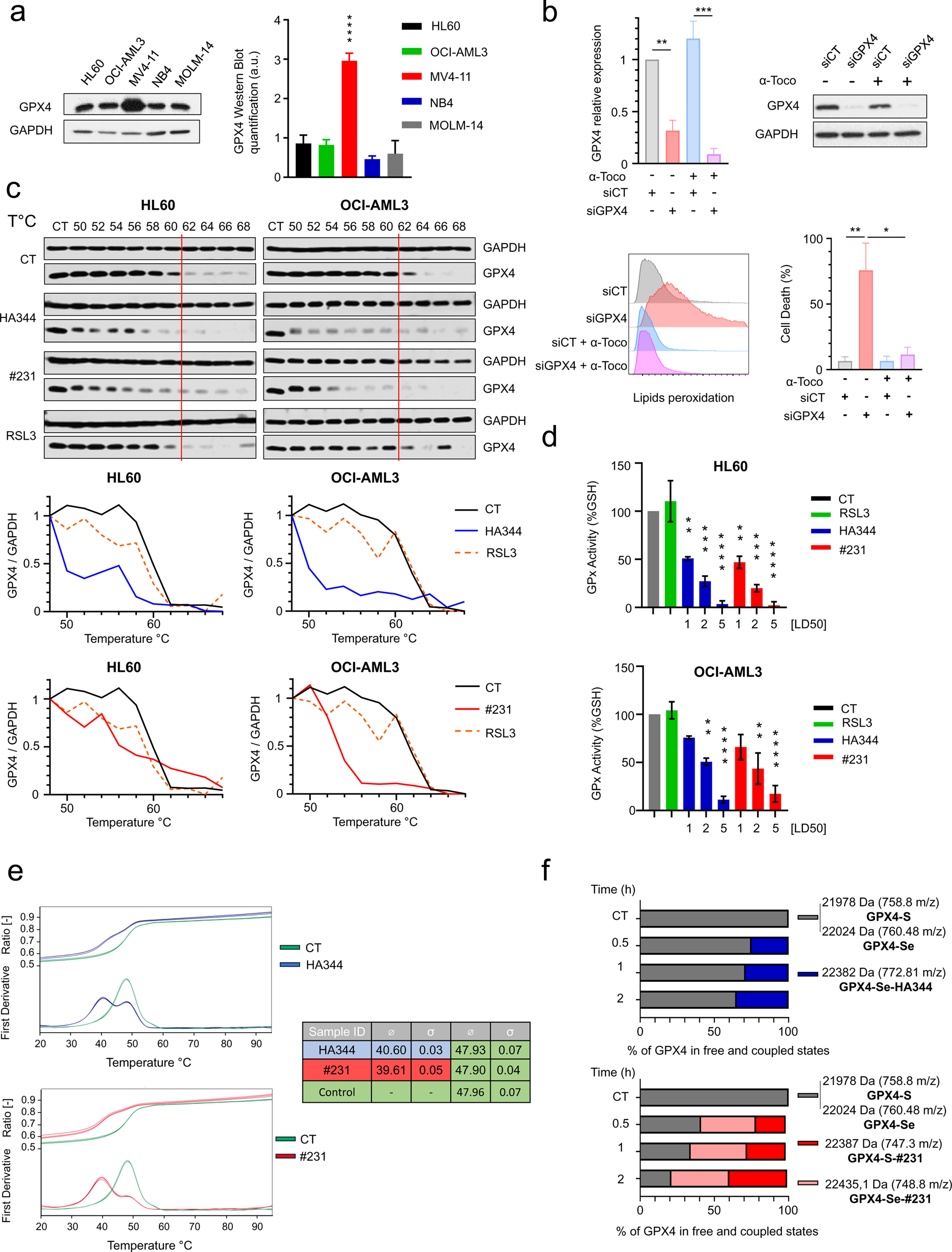
HA344 and #231 directly interact with GPX4, triggering ferroptosis. (a) GPX4 protein expression levels were determined by Western Blot analysis of HL60, OCI-AML3, MV4-11, NB4, and MOLM-14 cell lines (left panel). Subsequent quantification was realized with ImageJ software (right panel). **(b)** GPX4 expression was abolished in OCI-AML3 cell line through GPX4 siRNA transfection in the presence and absence of 1µM alpha tocopherol. The experiment was conducted over a 48-hour period, and GPX4 protein expression was monitored using Western Blot analysis (upper right panel). Additionally, relative RNA expression was determined using quantitative RT-PCR (upper left panel). Peroxidized lipids were assessed using the Lipid peroxidation assay (lower left panel), cell death induction was measured by DAPI incorporation and analyzed by flow cytometry (lower right panel). **(c)** Thermal Shift assay (CETSA) to analyze the interaction between HA344, #231, or RSL3 and GPX4. Cells were treated with 20 LD50 of HA344 and #231 or 40nM of RSL3 for one hour. The temperature was gradually increased with 2°C increments to disrupt the stability of the endogenous GPX4. Then, the different samples were subjected to Western blot analysis using an anti-GPX4 antibody (upper panel). Quantification of GPX4 protein was carried out with ImageJ software (right panel). **(d)** To assess GPX4 activity, we measured intracellular Reduced Glutathione (GSH) percentages levels in AML cells exposed to various concentrations of HA344 or #231, we used the GSH/GSSG Glo Assay kit and measured the resulting fluorescence signals with a plate reader. RSL3 was used as an inhibitor of GPX4 in this experiment **(e)** NanoDSF assay: Recombinant GPX4 samples at a concentration of 10 µM incubated in 2.5 % DMSO or with either 100µM HA344 or #231 for 2 hours at 21°C were analyzed. Measured denaturation curves in the temperature range from 25°C to 95°C and their corresponding 1st derivatives are presented. The average and standard deviations of the melting temperatures (Tm) obtained from the three technical replicates for each condition are depicted in the table. **(f)** Comprehensive Mass analysis for the identification of covalent complexes: a mixture of recombinant GPX4-S and GPX4-Se (ratio: 60/40) at a concentration of 10 µM was incubated with 100 µM HA344 and #231 for 30 minutes, 1 hour, and 2 hours at room temperature. The resulting product from this incubation was subjected to mass spectrometry analysis to identify the formed complexes.

To confirm a direct interaction of our compounds with GPX4, we performed thermal shift assays in the HL60 and OCI-AML3 cell lines **(Fig. 6c)**. One hour after HA344 or #231 addition, a significant GPX4 shift was observed compared to untreated condition. The modification of GPX4 migration likely reflected a post-translational modification or a major GPX4 conformational change induced by HA344 or #231. HA344 and #231 induced a drastic reduction in the melting temperature, which decreased from 62°C (under control conditions) to 56°C and 54°C for HA344 and #231 respectively in HL60 cells **(Fig. 6c, left panel)**. In OCI-AML3 cells, the impact of HA344 and #231 compounds was even higher, resulting in a decrease from 62°C to 50°C and 54°C respectively **(Fig. 6c, right panel)**. RSL3 was previously reported as a direct inhibitor of GPX4 ^21,27–29^. Surprisingly, in our hands, RSL3 failed to influence the melting temperature of GPX4, indicating that the interaction zone with GPX4 is different from that of our compounds or that RSL3 is not a direct GPX4 inhibitor but rather of TXNRD1, as recently described ^30,31^. Quantification of the western blot bands confirmed the lower stability of GPX4 when bound to HA344 or #231 **(Fig. 6c lower panels)**. We then treated HL60 and OCI-AML3 cell lines for 48h with increasing concentrations of HA344 and #231. Both compounds induced a dose-dependent reduction in GPX activity as assessed by the decrease in GSH level **(Fig. 6d)**. Intriguingly, RSL3, a previously described GPX4 inhibitor failed to reduce intracellular GSH levels in the same conditions.

All together our findings provide compelling evidence that both HA344 and #231 affect GPX4 stability **(Fig. 6c)**, leading to a drastic inhibition of expression and enzymatic activity, and to the concomitant collapse of the GSH pool **(Fig. 6d)**. Our results also highlight very different mechanisms of action between our compounds and RSL3, which again raises questions about RSL3’s ability to target GPX4 directly **(Fig. 6c)**. The recombinant GPX4 protein was synthesized following the procedure outlined by Borchert et al.^32^, featuring a selenocysteine in the active site. With this protocol, we can expect around 50% of the GPX4 protein that contained a selenocysteine. The effects of HA344 and #231 compounds on the recombinant GPX4 protein were evaluated using nano differential scanning fluorimetry. The GPX4 negative control exhibited a melting temperature (Tm) of 47.96 ± 0.07 °C. Upon treatment with 100 μM of HA344 or #231, GPX4 displayed a significant shift in Tm from 47.93°C/47.90°C to 40.60°C/39.61°C, respectively **(Fig. 6e, upper and lower panel)**. Ultimately, we assessed the binding ability of our compounds to the recombinant GPX4 protein through mass spectrometry. For the recombinant GPX4 protein alone, we obtained a mixture consisting of two forms, with and without selenium, according to a 40/60% ratio. After incubation for different times with the compounds, it was observed that HA344 bound to the GPX4-Se form but not to GPX4-S after 120 min, while #231 bound 100% of GPX4-Se within 30 minutes, and 40% of GPX4-S after 120 min. These data indicate that HA344 and #231 both bound GPX4-Se, but that #231 exhibited a significantly higher reactivity **(Fig. 6f).** Because of the strong affinity of our compounds for GPX4-Se, we carried out a click-chemistry experiment to investigate their binding to another selenocysteine-containing protein, namely TXNRD1. We confirm that HA344 and #231 can effectively bind to TXNRD1 in all three AML cell lines **(Fig. S7b)**, a finding corroborated by MS analysis (data not shown). Furthermore, both HA344 and #231 were found to inhibit TXNRD1 activity in a dose-dependent manner as efficiently as TRi-1 a well-characterized inhibitor of TXNRD1 ^30^ **(Fig. S7c)**. Additionally, unlike GPX4, the expression of TXNRD1 remains relatively consistent across the various cell lines **(Fig. S7d).**

## DISCUSSION

Under physiological circumstances, the generation of ROS constitutes a significant threat to cellular integrity. ROS can act on proteins, lipids, carbohydrates, and DNA, causing oxidative damage. Faced to these deleterious effects of ROS, the cell must promptly respond by initiating an antioxidant program aimed at inhibiting the various components responsible for ROS production and mitigating their impact on cellular function. The central player in the response to oxidative stress (OS) is the transcription factor NRF2 ^33^. In physiological conditions, NRF2 forms a complex with Keap1 leading to its ubiquitination by the E3 ubiquitin ligase Cullin 3 and subsequent degradation via the proteasome. However, when OS occurs, NRF2 is released from this cytoplasmic complex and relocates to the nucleus. To function as a transcription factor, NRF2 necessitates heterodimerization with the small MAF protein, enabling it to bind to specific consensus sequences known as antioxidant response elements (ARE) ^34–37^. ARE responsive genes assume crucial roles in safeguarding the cell through the production and restoration of GSH and TXN, regeneration of NADPH, management of heme and iron metabolism, and detoxification of ROS. GPX4 is crucial for cellular defense against OS and lipid peroxidation ^20^. GPX4 plays a key role in maintaining the integrity of cell membranes by catalyzing the reduction of lipid hydroperoxides into their alcohol counterparts, preventing the propagation of oxidative damage and ensuring cell survival. In addition, GPX4 dysregulation has been implicated in numerous pathological conditions, making GPX4 a critical player in cellular health and disease ^38^.

In the present study, we used a structure-activity analysis to identify small molecule inhibitor bearing a nucleoside and non-nucleoside core structure and exhibiting a potent anti-leukemic effect on different hematopoietic cell lines. Two lead compounds, HA344 and #231 showed very strong efficacy not only on a panel of AML cell lines but also in various bone marrow samples from AML patients, within the micromolar concentration range. HA344 and #231-mediated leukemic cell death was dependent on both apoptosis and ferroptosis, since both compounds activated caspase 3, 7 and 9 and induced lipid peroxidation whatever the AML cell lines tested. Additional studies, using a pharmacological approach confirmed the dependence of HA344 and #231 effect on apoptosis and ferroptosis. RNAseq analysis performed on different AML cell lines identified OS and apoptosis pathways as major determinants of the cell death programs triggered by our compounds. We also confirmed that both compounds induced nuclear relocation of NRF2 and the subsequent upregulation of ARE responsive genes as a defense mechanism.

Using click-chemistry *in cellulo* coupled to mass spectrometry we discovered among the putative HA344 and #231 targets some selenoproteins and more particularly GPX4. We next performed nanoDSF *in vitro* and thermal shift assay *in cellulo* and demonstrated that HA344 and #231 covalently bind to the selenocysteine (Se) present in GPX4 active site. This interaction resulted in the destabilization of the protein and its rapid degradation by an unknown mechanism. Very recent results in the literature have highlighted that ferroptosis inducers, including ML-162 and RSL3, originally believed to directly target GPX4, inhibited thioredoxin reductase 1 (TXNRD1) and that most of the observed effects of these drugs were mediated by TXNRD1 inhibition ^30^. During the screening process to identify FDA approval small molecules that target GPX4, lusutrombopag (a thrombopoietin agonist), navitoclax and venetoclax (two BH3-mimetics) were also discovered as potential GPX4 inhibitors ^39^. Whether these drugs frequently used in the clinic to treat AML patients inhibit GPX4 activity and whether they induce ferroptosis is currently unknown.

While our two compounds induced apoptosis and ferroptosis in all the leukemic cell lines tested, there are, however, some slight differences in their respective effects in different AML cell lines. Experiments using a mixture of recombinant GPX4-Se and GPX4-S protein allow to characterize the binding of both compounds to the enzyme. They highlighted that HA344 is less reactive to the enzyme but with a higher selectivity for GPX4-Se, while #231 is more reactive to the enzyme but with less selectivity for GPX4-Se as coupling to GPX4-S was also detected and showed an increase with incubation time.

As TXNRD1 was also significatively enriched among the potential target of our compounds, we performed click-chemistry experiments on several AML cell lines and confirmed a direct binding of HA-344 and #231 on TXNRD1. We also checked that both compounds were able to efficiently inhibit TXNRD1 activity in AML. In conclusion, HA344 and #231 exerts a dual activity on both GPX4 and TXNRD1 that could explain their very strong activity in AML cell lines.

Our findings may also have an impact on AML patient management in the future. Indeed, compelling studies have highlighted an increase of intracellular ROS both in cell lines and in leukemic cells derived from patients’ samples ^40–44^. Keeping an elevated intracellular ROS level within myeloid cells presents an important threat due to their high toxicity, requiring the establishment of a set of detoxification mechanisms to support their continued growth and viability. The use of compounds capable of selectively interfering with these detoxification processes will certainly help eliminating leukemia cells while minimizing the impact on healthy cells.

In conclusion, we describe here two new compounds capable of inducing both apoptosis and ferroptosis in different AML cell lines and CD34+ cells from AML patients. These compounds which exhibit high affinity for GPX4 and TXNRD1 hold great promise in the context of AML, where high GPX4 expression is associated with a pejorative prognosis. Thus, HA344 and #231 might represent new weapons in the therapeutic armamentarium of antileukemic drugs, either individually or in combination with currently available treatments.

## Legend to figures

**Supplemental Figure 1.**
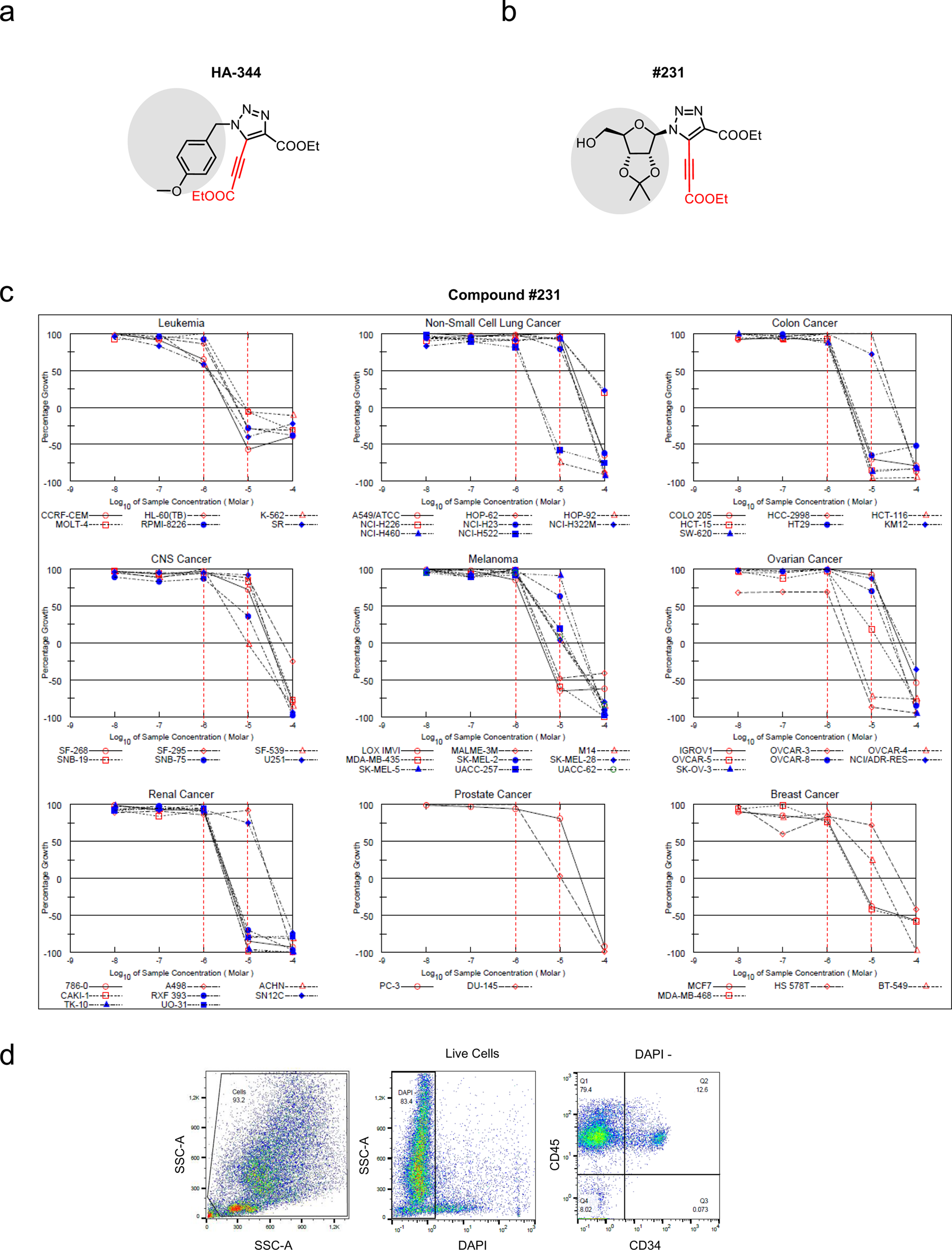
Representative structure of compounds **(a)** HA344 and **(b)** #231. **(c)** Efficacy of compound #231 in inducing cell death using the Human Tumor Cell Lines (NCI-60) platform of 60 different cancer cell lines. The dot plots presented in **(d)** represent the gating strategy employed for the analysis of cells derived from patients undergoing treatment. Living cells are identified based on their lack of Dapi incorporation, followed by the isolation of blast cells through positive labeling with CD45 and CD34.

**Supplemental Figure 2.**
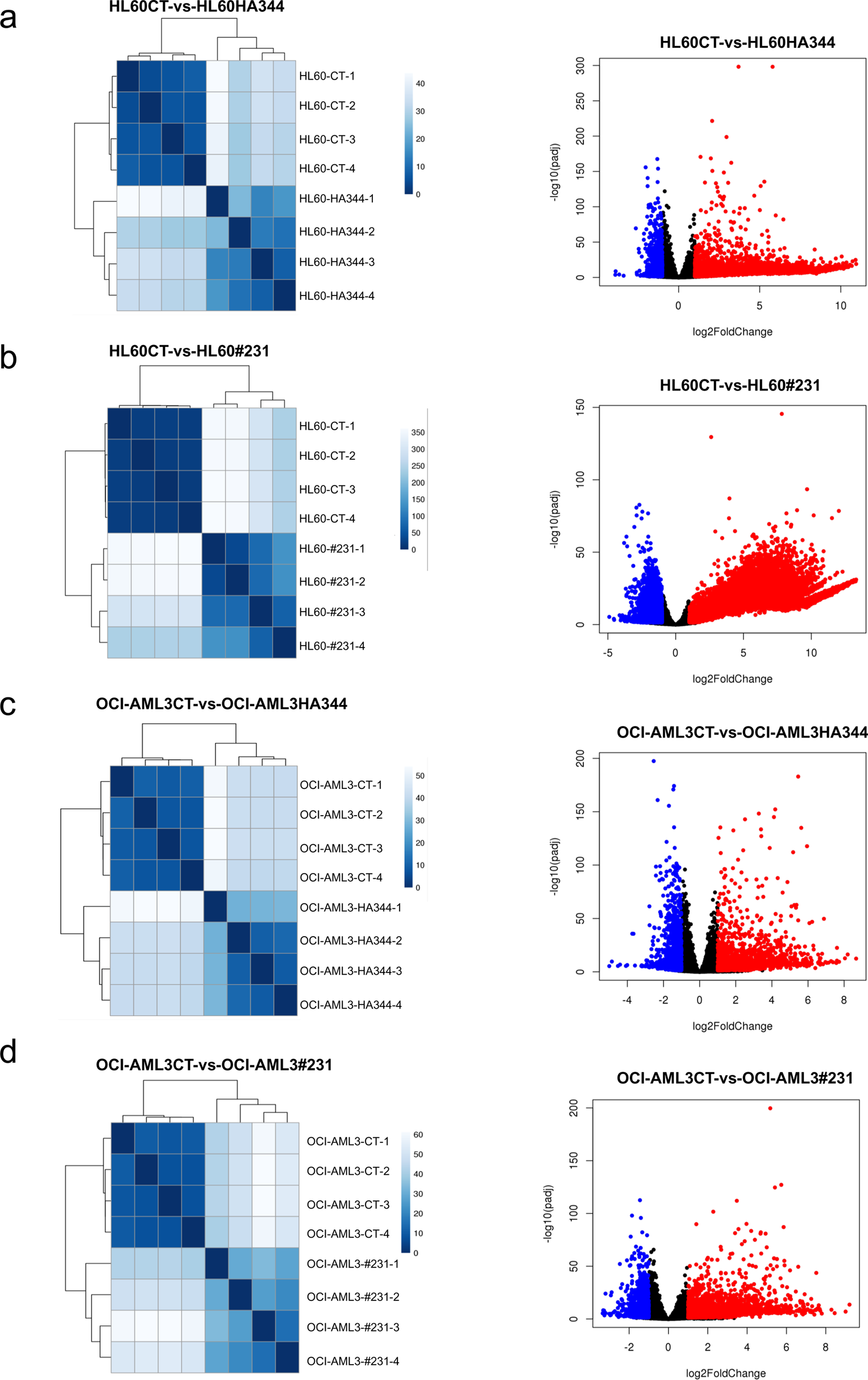
shows a Visualization of the sample distances for both control and treated conditions (left panel). Volcano plot representing genes that are upregulated and downregulated, expressed in log2 fold change within the same condition (right panel). These analyses are presented for the following pairs of conditions: **(a)** HL-60 CT vs HA344**, (b)** HL-60 CT vs #231**, (c)** OCI-AML3 CT vs HA344, and **(d)** OCI-AML3 CT vs #231.

**Supplementary Figure 3.**
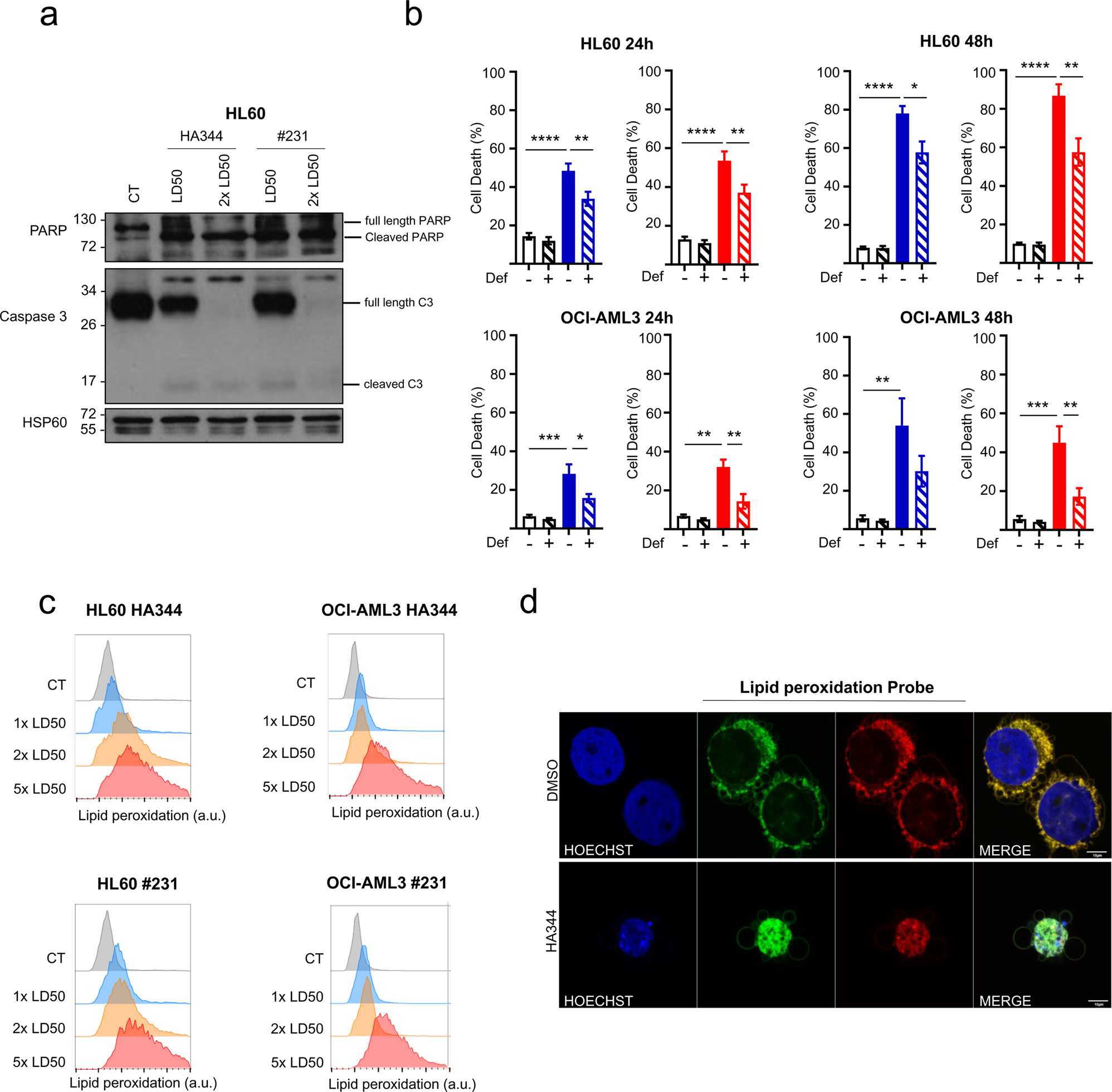
(a) HL60 cells were treated 24h by HA344 or #231 at LD50 or 2 LD50, then PARP and Caspase 3 cleavages were assessed by Western Blot. **(b)** HL60 and OCI-AML3 were pretreated with 5 µM deferoxamine followed by incubation with HA344 or #231 at their respective LD50 concentrations. Cell death was assessed at 24 and 48 hours by measuring DAPI incorporation using flow cytometry. **(c)** HL60 and OCI-AML3 cells were treated 3h with HA344 or #231 at 1, 2, and 5 LD50 concentrations, the quantification of peroxidized lipids was performed using a Lipid Peroxidation Assay Kit, followed by flow cytometry analysis. **(d)** HL60 cells were subjected to a 3-hour treatment with 5 µM of HA344. Lipid peroxidation induction was evaluated using a Lipid peroxidation assay kit. According to the transition from a 595/50nm (red) to 525/50nm (green) wavelength, alongside Hoechst nuclear staining, and subsequently observed through a confocal microscope imager.

**Supplementary Figure 4.**
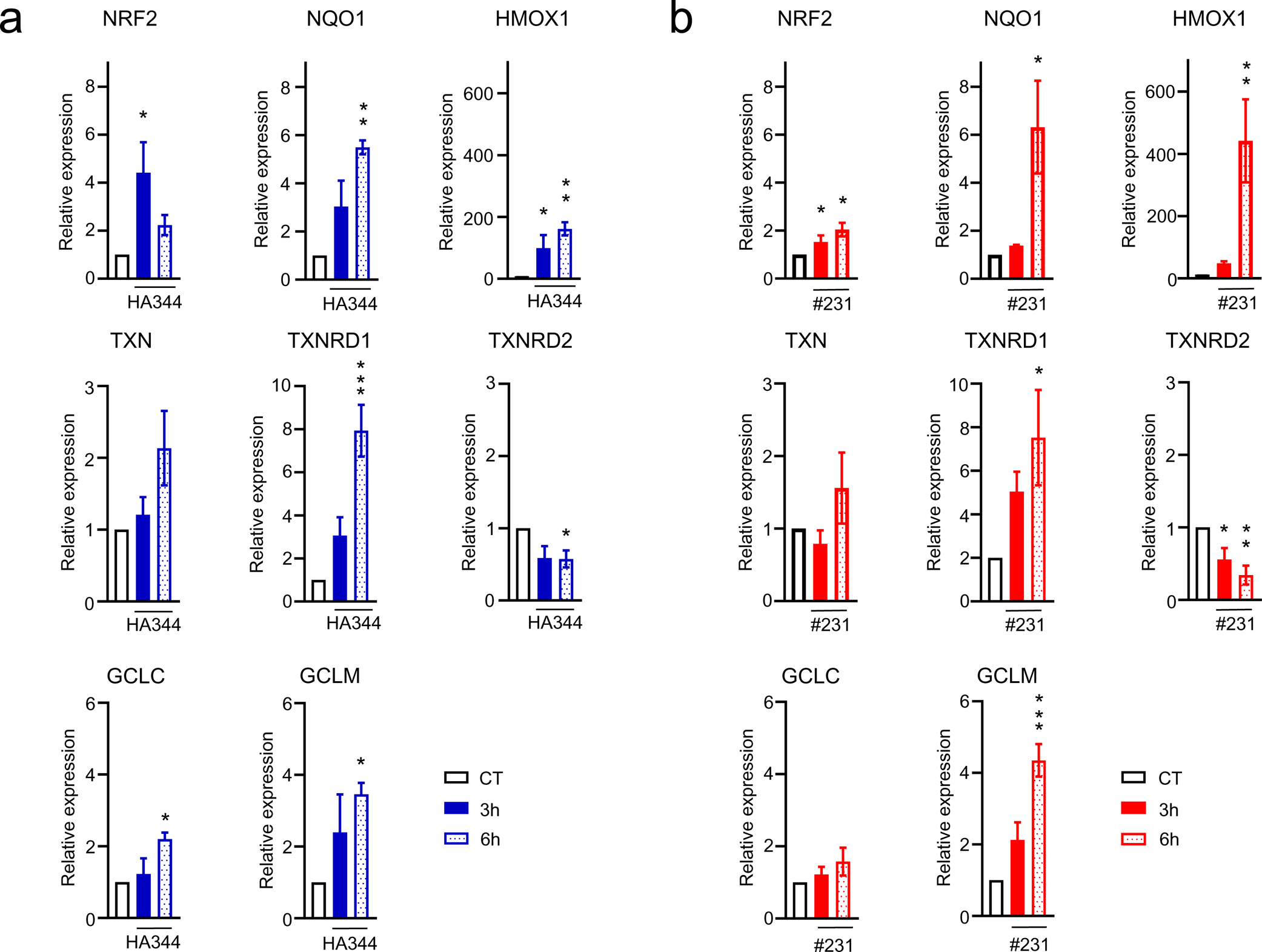
(a-b) Determining the mRNA expression levels of NRF2-dependent genes using RT-qPCR in OCI-AML3 cells after treatment with HA344 or #231 at LD50 for either 3 hours or 6 hours.

**Supplementary Figure 5.**
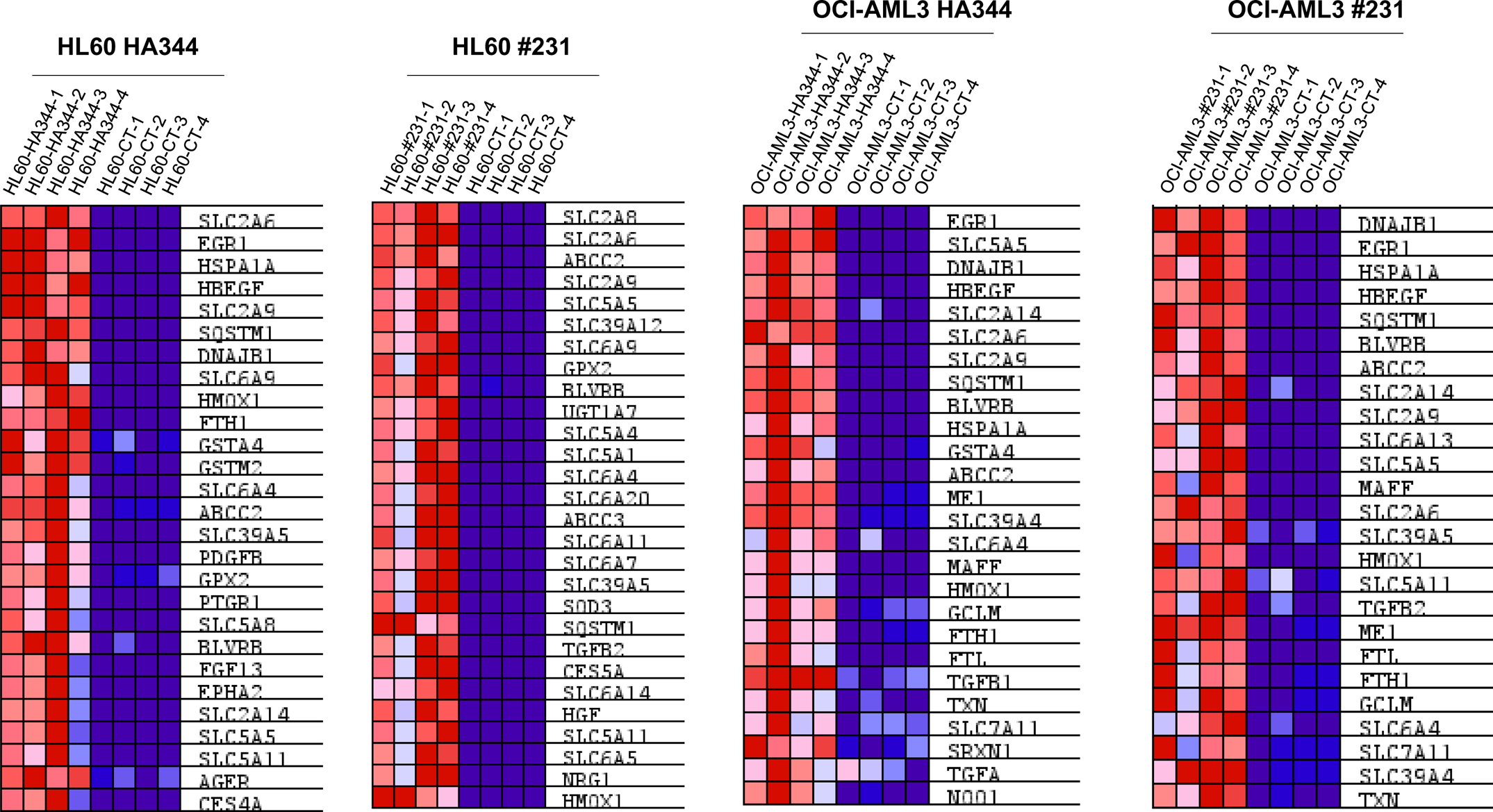
List of upregulated genes within the NRF2 pathway in HL-60 and OCI-AML3 cells treated with HA344 and #231.

**Supplementary Figure 6.**
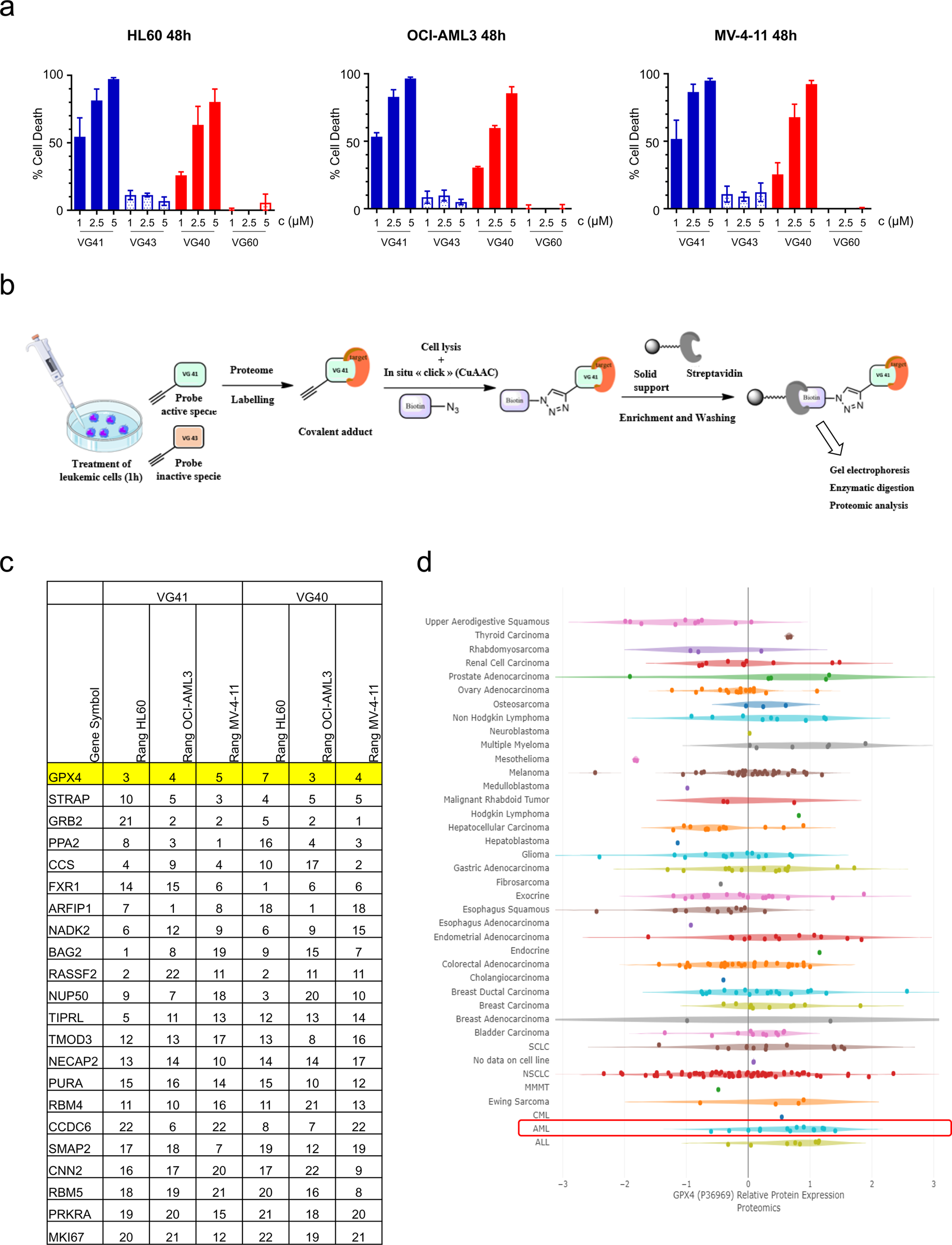
(a) Assessment of cell mortality resulting from the exposure of HL60, OCI-AML3, and MV4-11 cell lines to VG41, VG43, VG40, and VG60 at varying concentrations of 1µM, 2.5µM, and 5µM over a 48-hour duration. Cell death is determined by evaluating DAPI incorporation using flow cytometry. **(b)** visual illustration depicting the procedure employed for the Click Chemistry experiments. **(c)** Table highlighting the 22 proteins that are common to both our compounds and the three cell lines: HL60, OCI-AML3, and MV4-11. **(d)** Graph illustrating the protein expression levels of GPX4 in cells derived from patients with various cancer types. Data sourced from the DeepMap website.

**Supplementary Figure 7.**
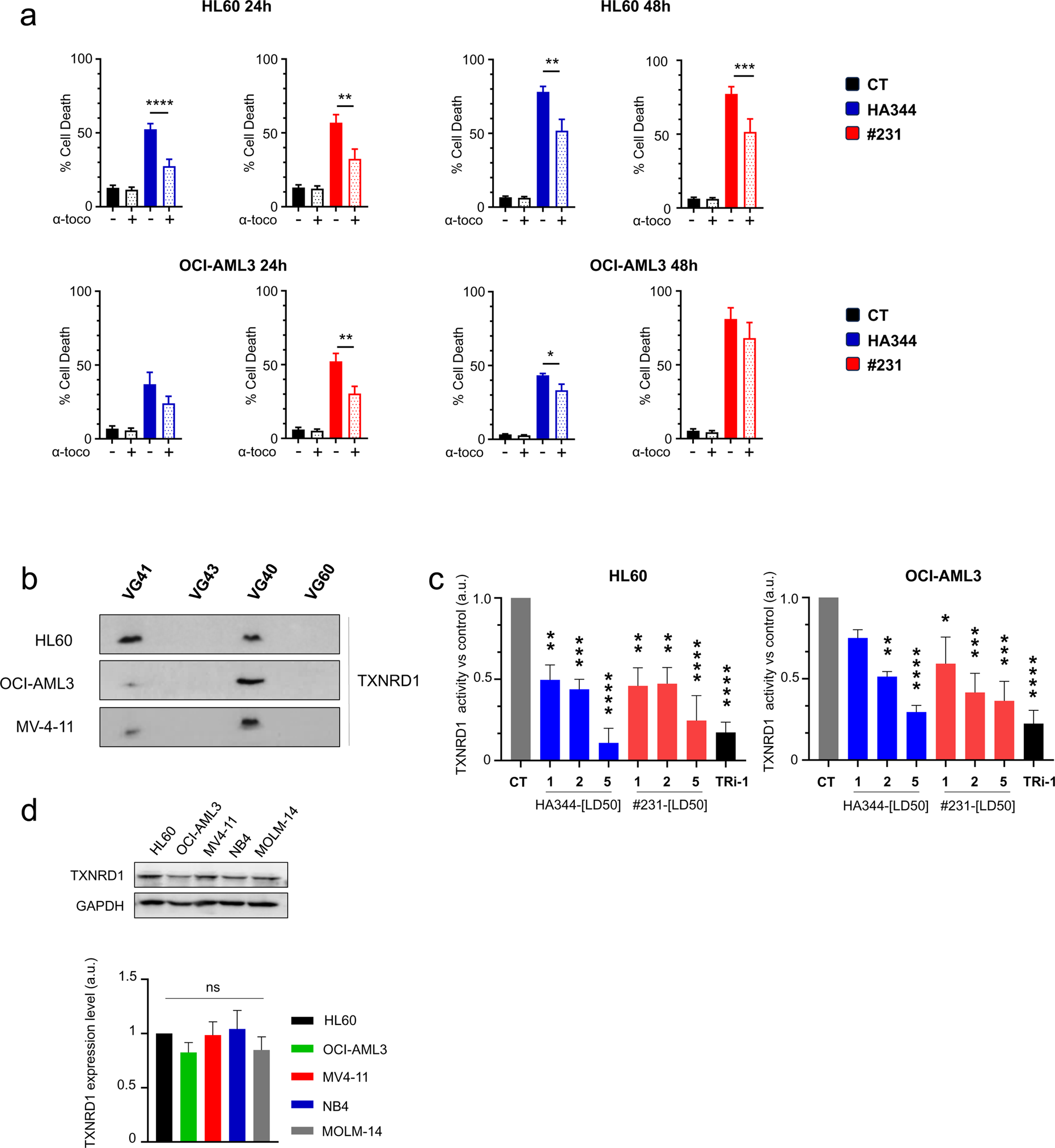
(a) The HL60 and OCI-AML3 cell lines were pretreated 1 hours with 1µM Alpha tocopherol before the addition of HA344 or #231 at their individual LD50 concentration during 24 and 48 hours. Cell death was quantified by measuring Dapi incorporation and conducting flow cytometry analysis. **(b)** HL60, OCI-AML3, MV4-11 cell lines, were subjected to treatment with 2.5 µM of VG41, VG43, VG40, and VG60 for 1 hour, followed by a click chemistry reaction and Western Blot analysis to confirm the binding of HA344 and #231 to TXNRD1. **(c)** To evaluate TXNRD1 activity, we quantified the reduction of DTNB to TNB using NADPH in AML cells treated with different concentrations of HA344 or #231. We employed the Thioredoxin Reductase Assay Kit and measured the resulting colorimetric signals at 412 nm using a plate reader. Tri-1 served as a specific inhibitor of TXNRD1 in this assay. **(d)** TXNRD1 protein expression levels were determined by Western Blot analysis of HL60, OCI-AML3, MV4-11, NB4, and MOLM-14 cell lines (upper panel). Subsequent quantification was realized with ImageJ software (lower panel).

**Supplemental Table 1.** Illustration of the genetic characteristics of the 25 patient BM samples used in this study, stratified on their genetic risk status at the time of initial diagnosis and classified as Intermediate or Adverse according to the European LeukemiaNet 2022 criteria ^45^. Detailed information includes their specific disease type, genotype, karyotype abnormalities, and the total number of treatment cycles they underwent. BM blast counts were determined using CD45, CD33, and CD34 labeling, and assessed through flow cytometry. Additionally, the percentage of blasts was cross-referenced with clinical histology laboratory findings, with only samples showing consistent results included in the study.

Intermediate Patients
| # Patient | Genotype- Karyotype | % blast cells |
| --- | --- | --- |
| 204 | AML - 46, XY [20] – 27 Azacytidine cycle | 20% |
| 204 | AML - 46, XY [20] - 39 Azacytidine cycle | 13% |
| 224 | AML - 46,XX[22] - DNMT3A(R882C) NPM1(W288CfsTer12) and PTPN11 (D106A, E69K, F71L and E76G) FLT3 ( Y591D) | 84% |
| 225 | AML - 46,XX[22], FLT3(FLT3-TKD : I836del), NPM1 (W288CfsTer12), PTPN11 (F71L, A72T, A72V et G503V), DNMT3A (G532V), FLT3 ( FLT3-TKD : A680V) and PRPF8(K1611N) | 80% |
| 236 | AML – 46,XY, NPM1 (W288CfsTer12), CEBP alpha mutation and FLT3 ITD , KRAS(G12A), TET2(N695NfsTer17), TET2(N1558PfsTer13), JAK2 (G571S) | 90% |
| 245 | AML - 46,XX[21] | 90% |
| 248 | MDS - 46,XX[20] | 10% |
| 250 | MDS – 46,XY[21] | 9% |
| 253 | AML – 46,XX[21] | 35% |
| 254 | AML – 46,XY[20] | 96% |

Adverse Patients
| # Patient | Genotype- Karyotype | % blast cells |
| --- | --- | --- |
| 219 | AML - 46,X,t(X;11)(p14;q14)[9]/47,idem,+der(X)t(X;11)[12] ; 3 MLL genes | 51% |
| 222 | AML - 50,XX,+5,+8,+10,+13[4]/46,XX[16] - DNMT3A(R882C),NMP1(W288CfsTer12) FLT3-ITD | 80% |
| 227 | AML - 46-50,XY,-5,der(6)t(6;?)(p11;?),+8,-11,-11,-2-7mar[cp8]/46-49,XY,-5,der(6)t(6;?)(p11;?),-16,der(17)t(17;?)(p11;?)[cp7] | 83% |
| 228 | MDS - 44,X,-X,-3,der(4)t(3;4)(q11;q32),del(5)(q22q34),t(9;17)(p12;p12),-12,+mar[10] | 18% |
| 231 | AML - 47,XX,dup(3)(q21;qter),+8[8]/47,XX,der(7)t(3;7)(q21;p22),+8[2], mutation pGTrpfs*12 of ASXL1, NPM1 (W288CfsTer12), SRSF2(P95R), KRAS(G13D), TET2(Q1051RfsTer4 et S1190LfsTer36), SH2B3 (R189Q). | 92% |
| 233 | AML - 46,XX,add(10)(p14)[22] | 87% |
| 238 | AML - 46,XX, inv(16)(p13q22)[6]/46,XX[17] fusion CBFβ::MYH11 | 40% |
| 239 | AML – 46,XY, IDH1 and IDH2 mutation, 2 ASXL1 mutations, 2 DNMT3A mutation, EZH2, RUNX1 and SRSF2 mutation | 20% |
| 240 | AML - 46,XX,del(5)(q32q34)[22] | 67% |
| 241 | AML - 47XX,t(4;15)(q26;23),+8,del(11)(q14),add(21)(q2?1)[5]/ 48,s1,+8[2]/ 49,sd11,+der(4)t(4;15),del(10)(q2?3)[1]/ 51,sd12,+13,+20[2]/ 46,XX[10] FLT3-, IDH1-,IDH2-,NPM1-, mutation of the exon 8 and 9 of TET2 and of exon 8 of CBL. | 80% |
| 243 | AML - 46,XX,del(13)(q13q21)[6]/46,XY,der(17)t(17;?)(q25;?)[3]/46,XY[18] | 95% |
| 246 | MDS - 46,XY,inv(3)(q21q26),r(7)[9]/46,XY[11] | 19% |
| 247 | AML - 44,X,-Y, dic(3;16)(3pter->3q10::?:16qter),del(5)(q15q34),-7,del(12)(q22),der(17)t(-Y;17)(q11;p12),del(21)t(21;?)(p12;?),r(21)[9]/46,XY[2] | 28% |
| 208 | MDS - TP53 (R158H and H214R), TET2(P1419H) - 46,XX,del(5)(q13q32)[2]/46,XX[8] – 13 Azacytidine Venetoclax cycle + 2 Azacytidine cycle | 8% |
| 257 | AML - 46,XX,del(9)(q12q32)[3]/46,XX[12] | 65% |

**Supplementary Table 2.**
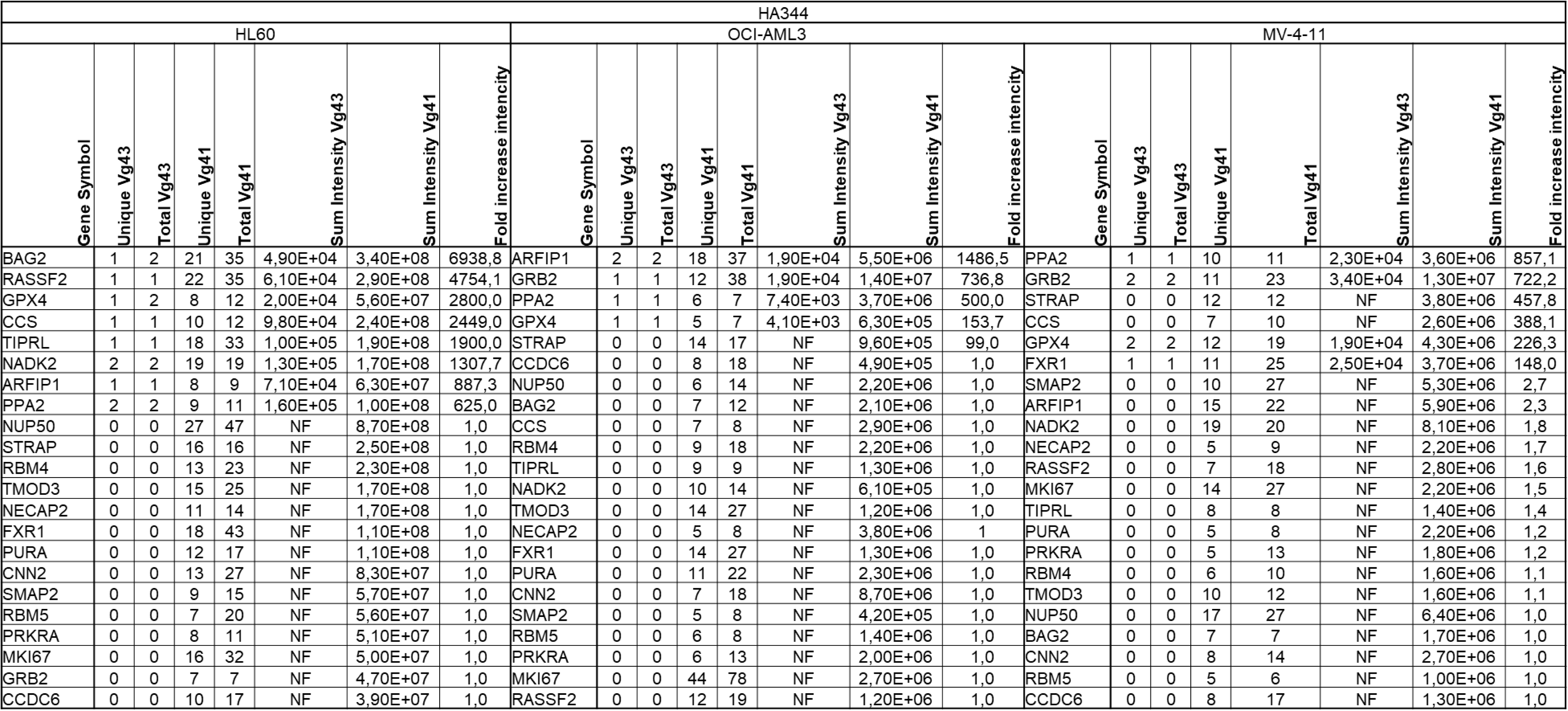
A summary table presenting 22 potential target proteins shared among various cell lines (HL60, OCI-AML3, and MV4-11) treated with HA344 identified by Mass spectrometry following Click chemistry experiment.

**Supplementary Table 3.**
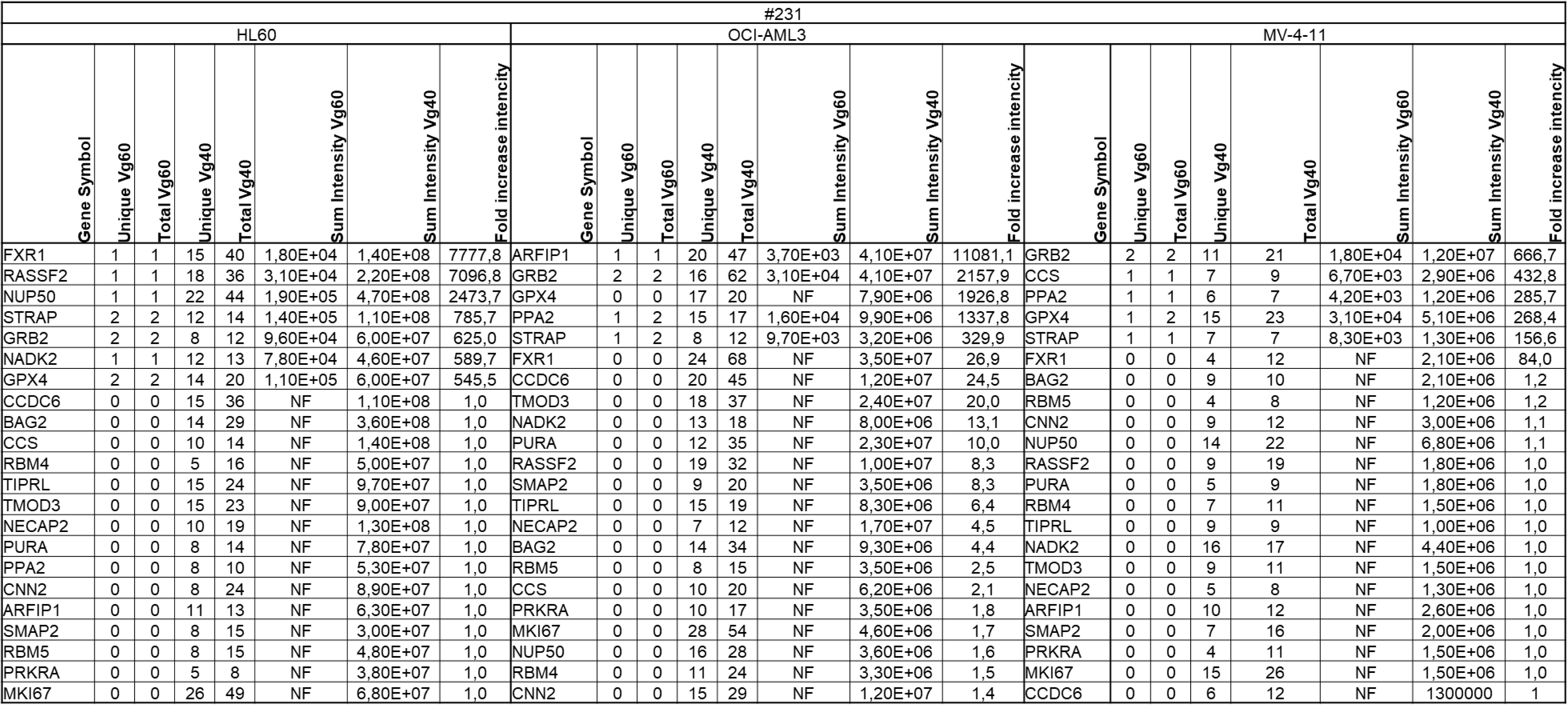
A summary table presenting 22 potential target proteins shared among various cell lines (HL60, OCI-AML3, and MV4-11) treated with #231 identified by Mass spectrometry following Click chemistry experiment.

## Supporting information

Chemical Synthesis

## Acknowledgements

This research was supported by the INSERM, Université Côte d’Azur, the Fondation ARC pour la Recherche contre le Cancer (Equipe Labellisée 2022-2025), the Canceropole PACA, a grant from PLBio (R19003AP-2019-2023), ITMO cancer 2019-2023 and the Association Laurette Fugain (ALF). This work was also funded by the French government (National Research Agency, ANR) through the “Investissement for the future” LABEX SIGNALIFE program reference #ANR-11-LABEX-0028-01. C Favreau has been a recipient from a fellowship from PLBio and is currently recipient of a grant from the Institut Carnot OPALE. This work was supported by the French Infrastructure for Integrated Structural Biology (FRISBI) ANR-10-INBS-0005. We thank Luc Negroni (IGBMC proteomics platform) for mass spectrometry analysis of recombinant GPX4 protein. We sincerely thank the GIS-IBISA multi-sites platform Microscopy Imagery Côte d’Azur (MICA), and particularly the imaging site of C3M (INSERM U1065) supported by Conseil Régional, Conseil Départemental, and IBISA. The help of Marie Irondelle is acknowledged.

## Author contributions

C.F., C.S., M.B., T.B., S.B., M.Z., E.K.: Acquisition of data, conception of figures. F.G., C.B., A.P. Acquiring and analyzing experimental data concerning the recombinant protein GPX4. A.V., J.D.G. Synthesis and characterization of chemical molecules. M.L.A, T.C., S.R., A.J.:Authors discussed the results and commented on the manuscript. R.B. P.A., AR.M.:Editing the article, analysis and interpretation of results. G.R:Acquisition of data, drafting the article, analysis and interpretation of relevant studies, conception of the figures.

## Material and Methods

All data relating to the protocols and materials used in this article are available in the online document entitled “material and methods”.

