## Supplementary material for "Dual targeting of GPX4 and TXNRD1 triggers eradication of AML cells through induction of apoptosis and ferroptosis": Material and Methods

### Chemical synthesis

HA344, VG41 and VG43 were prepared as reported in Zerhouni *et al.*[23] . #231 was synthesized according to our prior report: Amdouni et al. J. Med. Chem., 2017, 60, 1523. VG40 and VG60 were synthesized as described in the "chemical synthesis" document.

### Cell lines

The human AML, HL60 (CCL-240) and MV4-11 (CRL-9591) cell lines were provided by ATCC (Manassas, VA, USA). The OCI-AML3 (ACC 582), NB4 (ACC 207) and MOLM-14 (ACC 777) cell lines were provided by DSMZ (Leibniz, Germany), and were grown in RPMI 1630 medium (Lonza, Walkersville, MD, USA), all media were supplemented with 10% of FBS, 1% of pyruvate sodium. All cell cultures were grown at 37°C under 5% CO2, 50 units/ml penicillin and 50 mg/ml streptomycin to minimize contamination.

### PBMC

Human peripheral volunteers were obtained from healthy donors with informed consent following the Declaration of Helsinki according to recommendations of an independent scientific review board. The project has been validated by The Etablissement Français du Sang, the French national agency for blood collection (13-PP-11). Blood samples were collected using ethylene diamine tetra acetic acid–containing tubes. Mononucleated cells were first isolated using Ficoll Hypaque (Eurobio, CMSMSL0101). PBMC were grown in RPMI 1640 medium with glutamax-I (Life Technologies, 61870044) supplemented with 10% (vol/vol) fetal bovine serum (Life Technologies) at 37°C under 5% CO2, 50 units/ml penicillin and 50 mg/ml streptomycin to minimize contamination.

### Bone Marrow samples preparation

Bone marrow samples are received and subjected to 10-minute hemolysis with 20 ml of lysis buffer (#555899, BD Biosciences). Cells were then washed with 30 ml of Dulbecco's Phosphate Buffered Saline (PBS, #14190-094, Gibco) and centrifuged for 5 minutes at 400g. After several washing steps, cells were counted with trypan blue solution (#T8154, Sigma). 200 000 bone marrow cells are resuspended in 100µl of running buffer and incubated for 10 min in the presence of 1µl of FcR Blocking Reagent (130-059-901, Miltenyi Biotec) at 4°C. cells were then stained with 1µl of anti-CD45 (#3047014, Biolegend), anti-CD33 (#36618, Biolegend), and anti-CD34 (#343510, Biolegend) antibodies. Cells were finally washed and resuspended in 200 µl of Running buffer. Subsequently, the samples were analyzed using a MacsQuant 16 cytometer (Miltenyi Biotec), Flowjo software was used for cytometry data analysis.

### Medium for Bone Marrow cells

Complete medium, used for bone marrow cell culture is composed of IMDM medium (LONZA) supplemented with 2% BIT, 20% SVF, 100 µM b-mercaptoethanol,10 ng/ml GCSF, 300 ng/mL SCF, 300 ng/mL FLT3L, and 10 ng/mL interleukin-6 (IL6).

### Antibodies

Caspase 3 (9668, RRID:AB_2069870), PARP (9542, RRID:AB_2160739), NRF2 (12721, RRID:AB_2715528, Lamin A/C (4777, RRID:AB_10545756),TRXR1 (15140, RRID:AB_2798725), anti-rabbit IgG HRP-linked (7074, RRID:AB_2099233) and anti-mouse IgG HRP-linked (7076, RRID:AB_330924) antibodies were purchased from Cell signaling Technology CST (Danvers, MA, USA). HSP90 (sc-13119, RRID:AB_675659) and Rho DGIa (sc-373724, RRID:AB_10917570) were purchased from Santa Cruz Biotechnology (Dallas, TX, USA). GPX4 (SAB4300725, AB_3075489) antibody was purchased from Sigma-Aldrich (Saint-Louis, MO, USA). GAPDH (ab9485, RRID:AB_307275) antibody was purchased from Abcam (Cambridge, UK).

### Reagents and Inhibitors

Q-VD-OPh (A1901-10mg) was purchase from Clinisciences (Nanterre, France). Ciclopirox (S2528) and TRi-1 (S0267) were purchase from Selleckchem (Cologne, Germany). Deferoxamine mesylate (5784) and RSL3 (6118) were purchase from Tocris (Bristol, UK). (+)α-Tocopherol (T3634) was purchase from sigma-aldrich (Saint-louis, MO, USA).

### Caspases assay

After treatment, cells were subjected to lysis for 30 min at 4°C using a lysis buffer composed of HEPES (50mM), NaCl (150mM), EDTA (20mM), and Triton (0.2%). Then for each sample, 25µg of extract was deposited in quadruplicate in a 96-well plate containing 250µM AMC substrate (DEVD for caspases 7 and 3 or IETD for caspase 9), supplemented with 10µM DTT. One of the quadruplicate point contained a specific inhibitor of each AMC substrate (DEVD-CHO or IETD-CHO) in order to assess the specific activity of each protease. Fluorescence readings were then taken for 1 hour and 30 minutes with a Synergy H1 plate reader (Bio Tek) at 390nm excitation and 460nm emission.

### Cellular thermal shift assay (CETSA)

CETSA experiments were performed using the protocol described by Jafari R. in 2014. Cells were first exposed for 1h to a high concentration of HA344 and #231 corresponding to 20 times the LD50. The cells were then divided into different conditions corresponding to the required temperature points. Each cell sample was subjected to a 3-minute heat treatment at the specified temperature, followed by a 3-minute cooling to 25°C, then mechanical lysis in liquid nitrogen. Proteins were extracted by centrifugation at 17,000g at 4°C for 10 minutes. A solution of 4x Laemmli buffer was added to each sample and heated for 5 minutes at 95°C. Treated samples were then analyzed by western blot.

### Click chemistry

The click chemistry experiments were conducted according to Chen Y-C, Zhang C. Genes & Cancer 2016. Briefly, cells were seeded and treated for 1hr with 2µM VG41 (clickable HA344 molecule provided by the ICN) or VG43 (negative control for the experiment), VG40 (clickable #231 molecule provided by the ICN) or VG60 (negative control for the experiment). After treatment, cells were washed three times in PBS and the pellet was resuspended in 1330 µL of 1 % NP40 buffer completed with protease inhibitor. Resuspended cells were sonicated for 30s three times and incubated 30 min on ice before centrifugated at 20 000 g for 10 min. After cell lysis, proteins were extracted and quantified using Bradford method. For each condition, up to 10mg of proteins in 9.4 ml of 1 % NP40 buffer was used to perform the biotin addition using 100 µL of biotin azo-azide at 5 mM, 200 µL TCEP at 50 mM, 100 µL TBTA at 10 mM and 200 µL $CuSO_4$ at 50 mM. Proteins were then incubated 1 hr in the dark at room temperature. After incubation, 40ml ice-cold methanol was added. Proteins in methanol were precipitated overnight at -20°C. Proteins were centrifugated at 5200 g, 30 min at 0°C and washed 3 times in the same conditions using ice-cold methanol. After drying, proteins were resuspended in the resuspension buffer (6 M urea, 2 M thiourea, 10 mM HEPES) and sonicated. 40 µL of DTT were added to the proteins which were then incubated 40 min at room temperature. 40 µL of iodoacetamide were added to the samples and incubated 30 min at room temperature in the dark. After pre-washing of the streptavidin beads, resuspended proteins were added to the beads in a 15 ml tube and incubated on a rotor for 2h at room temperature. Beads were collected by centrifugation at 200 g for 3 min at room temperature and then washed with resuspension buffer (PBS and 1 % SDS, PBS) solution twice. After washing, elution was performed using sodium dithionite solution. Eluted proteins were incubated overnight at -20°C with ice-cold methanol. Proteins were then pelleted and dried before being resuspended in 4 % SDS buffer and 2X SDS free loading buffer. Samples were then prepared for classical Western Blot.

### DCFDA

Intracellular ROS levels were measured by DCFDA fluorescence probe (Sigma, Saint-Louis, MO, USA). AML cell lines were incubated in presence of 10 µM DCFDA for 30 min at 37°C. Relative DCFDA fluorescence was measured using a flow cytometer. Results were expressed in arbitrary fluorescence units.

### GPx Activity

The measurement of GPX4 activity involves quantifying the intracellular level of GSH (GSH-GLo™ Glutathione assay kit V6912), following the guideline provided by Promega (Madison, WI, USA). Briefly, A GSH standard range is established for the quantification of our samples. Subsequently, 40 000 cells are suspended in 50 µl of PBS and duplicate-plated in a black plate. To each well, 1 µl of 50 mM TCEP is added and shaked for 5 minutes at room temperature (RT). Following this, 50 µl of GSH-Glo™ Reagent 2x is introduced to the well and shaked for 30 minutes at RT. Finally, 100 µl of Luciferin Detection Reagent is added, and the mixture is incubated for 15 minutes at RT. Luminescence is then measured using a Synergy H1 plate reader (Bio Tek, Winooski, VT, USA).

### TXNRD1 Activity

TXNRD1 activity was assessed by measuring the reduction of DTNB to TNB using NADPH, as per the protocol outlined by Merck (Rahway, NJ, USA). Briefly, AML cells were treated with 1, 2, or 5 DL50 of HA344 or #231 for 1 hour. Cell lysates (30 µg) were then incubated in a buffer containing NADPH (180 µg/ml) and DTNB (1.2 mg/ml). The conversion of DTNB to TNB was quantified by monitoring the colorimetric signal at 412 nm using a Synergy H1 plate reader (Bio Tek, Winooski, VT, USA). Tri-1 was employed as a specific inhibitor of TXNRD1 in this assay.

### Mass Spectrometry Analysis

Protein samples from click chemistry experiment were sent to Ross Tomaino at Taplin Biological Mass Spectrometry Facility (Harvard Medical School). Excised gel bands were cut into approximately 1 $mm^3$ pieces. The samples were reduced with 1 mM DTT for 30 mins at 60°C and then alkylated with 5 mM iodoacetamide for 15 min in the dark at room temperature. Gel pieces were then subjected to a modified in-gel trypsin digestion procedure. Gel pieces were washed and dehydrated with acetonitrile for 10 min followed by removal of acetonitrile. Pieces were then completely dried in a speed-vac. Rehydration of the gel pieces was with 50 mM ammonium bicarbonate solution containing 12.5 ng/µl modified sequencing-grade trypsin (Promega, Madison, WI) at 4°C. Samples were then placed in a 37°C room overnight. Peptides were later extracted by removing the ammonium bicarbonate solution, followed by one wash with a solution containing 50 % acetonitrile and 1 % formic acid. The extracts were then dried in a speed-vac (~1h). The samples were then stored at 4°C until analysis.

On the day of analysis, the samples were reconstituted in 5 -10 µl of HPLC solvent A (2.5 % acetonitrile, 0.1 % formic acid). A nano-scale reverse-phase HPLC capillary column was created by packing 2.6 µm C18 spherical silica beads into a fused silica capillary (100 µm inner diameter x ~30 cm length) with a flame-drawn tip. After equilibrating the column each sample was loaded via a Famos auto sampler (LC Packings, San Francisco CA) onto the column. A gradient was formed, and peptides were eluted with increasing concentrations of solvent B (97.5 % acetonitrile, 0.1% formic acid).

As each peptide was eluted, they were subjected to electrospray ionization and then they entered into an LTQ Orbitrap Velos Pro ion-trap mass spectrometer (Thermo Fisher Scientific, San Jose, CA). Eluting peptides were detected, isolated, and fragmented to produce a tandem mass spectrum of specific fragment ions for each peptide. Peptide sequences (and hence protein identity) were determined by matching protein or translated nucleotide databases with the acquired fragmentation pattern by the software program, Sequest (ThermoFinnigan, San Jose, CA). The modification of 79.9663 mass units to serine, threonine, and tyrosine was included in the database searches to determine phosphopeptides. Phosphorylation assignments were determined by the Ascore algorithm. All databases include a reversed version of all the sequences and the data was filtered to between a one and two percent peptide false discovery rate.

### Lipid peroxidation assay

Cells were treated with HA344 or #231, then loaded with 200µl Lipid Peroxidation Assay Sensor (Lipid peroxidation assay kit, cell-based, ab243377, Abcam) diluted 1:5000 in PBS. Cells were then incubated for 30 minutes at 37°C with 5% CO2 in the dark. cells were washed 3 times and finally resuspended in 200 µl PBS. Flow cytometry was then carried out (Ex 488 Em 530nm and 572nm) to assess lipid peroxidation levels, calculated by dividing the fluorescent median values of the 530nm channel by those of the 572nm channel.

### RNA-sequencing

#### RNA extraction

RNA was prepared from $6 \times 10^6$ cells using the RNeasy Mini Kit according to manufacturer's protocol (Qiagen, 74104).

#### RNA Library Preparation and NovaSeq Sequencing

RNA Library Preparation and NovaSeq Sequencing RNA samples were quantified using Qubit 4.0 Fluorometer (Life Technologies,Carlsbad, CA, USA) and RNA integrity was checked with RNA Kit on Agilent 5300 Fragment Analyzer (Agilent Technologies, Palo Alto, CA, USA). RNA sequencing libraries were prepared using the NEBNext Ultra RNA Library Prep Kit for Illumina following manufacturer's instructions (NEB, Ipswich, MA, USA). Briefly, mRNAs were first enriched with Oligo(dT) beads. Enriched mRNAs were fragmented for 15 minutes at 94 °C. First strand and second strand cDNAs were subsequently synthesized. cDNA fragments were end repaired and adenylated at 3'ends, and universal adapters were ligated to cDNA fragments, followed by index addition and library enrichment by limited-cycle PCR. Sequencing libraries were validated using NGS Kit on the Agilent 5300 Fragment Analyzer (Agilent Technologies, Palo Alto, CA, USA), and quantified by using Qubit 4.0 Fluorometer (Invitrogen, Carlsbad, CA).

The sequencing libraries were multiplexed and loaded on the flowcell on the Illumina NovaSeq 6000 instrument according to manufacturer's instructions. The samples were sequenced using a 2x150 Pair-End (PE) configuration v1.5. Image analysis and base calling were conducted by the NovaSeq Control Software v1.7 on the NovaSeq instrument. Raw sequence data (.bcl files) generated from Illumina NovaSeq was converted into fastq files and de-multiplexed using

Illumina bcl2fastq program version 2.20. One mismatch was allowed for index sequence identification.

**Data Analysis**

After investigating the quality of the raw data, sequence reads were trimmed to remove possible adapter sequences and nucleotides with poor quality using Trimmomatic v.0.36. The trimmed reads were mapped to the Homo sapiens reference genome available on ENSEMBL using the STAR aligner v.2.5.2b. The STAR aligner is a splice aligner that detects splice junctions and incorporates them to help align the entire read sequences. BAM files were generated as a result of this step. Unique gene hit counts were calculated by using feature Counts from the Subread package v.1.5.2. Only unique reads that fell within exon regions were counted. After extraction of gene hit counts, the gene hit counts table was used for downstream differential expression analysis. Using DESeq2, a comparison of gene expression between the groups of samples was performed. The Wald test was used to generate pvalues and Log2 fold changes. Genes with adjusted p-values < 0.05 and absolute log2 fold changes > 1 were called as differentially expressed genes for each comparison. A gene ontology analysis was performed on the statistically significant set of genes by implementing the software GeneSCF. The goa_human GO list was used to cluster the set of genes based on their biological process and determine their statistical significance. A PCA analysis was performed using the "plotPCA" function within the DESeq2 R package. The plot shows the samples in a 2D plane spanned by their first two principal components. The top 500 genes, selected by highest row variance, were used to generate the plot.

-**Overall Sequencing Statistics :**

| | Total Reads | Total Mapped Reads | % Total Mapped Reads | Unique Mapped Reads | % Unique Mapped Reads |
|---|---|---|---|---|---|
| Mean | 36369092 | 35293229 | 97 | 33421208 | 92 |
| SD | 7762627 | 7440938 | 2 | 7413615 | 3 |

### Real Time-quantitative PCR (RT-qPCR)

RNA was prepared from $5.10^6$ cells using the RNeasy Mini Kit according to manufacturer's protocol (Qiagen, 74104). Each cDNA sample was prepared using AMV RT and random primers (Promega, M510F and C1181). Real-time polymerase chain reaction (PCR) was performed using the SyBR Green detection protocol (Life Technologies, 4367659). Briefly, 5 ng of total cDNA, 500 nM (each) primers, and 5 µL SyBR Green mixture were used in a total volume of 10 µL. Detection of endogenous control L32 was used to normalize the results. Specific forward and reverse primers are accessible upon request.

### Subcellular fractionation

Cells were washed with wash buffer (proteoExtract Subcellular Proteome Extraction KIT, Calbiochem, La Jolla, CA, USA). Then, cells were pelleted by centrifugation (10 min at 300 g).

The wash buffer was aspirated and discarded and Extraction Buffer I was added to extract cytosolic fraction. Cells were incubated for 10min at 4 °C under gentle agitation, centrifugated for 10 min at 800g. The supernatant contains the cytoplasmic fraction. Extraction Buffer II was added to the pellet and cells were incubated for 30 min at 4°C under gentle agitation. Then, cells were centrifugated for 10 min at 5500xg and the supernatant containing the microsomal fraction was collected. This crude microsomal fraction contains the plasma membrane, mitochondria, endoplasmic reticulum, Golgi apparatus and lysosomes. Extraction Buffer III was added to the pellet and cells were incubated for 30 min at 4°C under gentle agitation. Then, cells were centrifugated for 10 min at 6800xg and the supernatant containing the nuclear fraction was collected. A total of 30 µg of cytosolic and nuclear fractions were separated on 15% polyacrylamide gel and transferred onto PVDF membrane (Chemicon, Millipore, Billerica, MA, USA). After blocking non-specific-binding sites in saturation buffer, the membranes were incubated with anti-NRF2 antibody. Rho GDI and Lamin A/C were used as control of cytosolic and nuclear fractions respectively.

#### Electroporation of leukemia cells

OCI-AML3 cells were transfected by electroporation at various conditions using the electroporation system Amaxa (Lonza, Basel, Switzerland). In brief, $2.10^6$ cells were suspended in 100 µl complete Amaxa buffer in electroporation cuvette, 300 nM of control or GPX4 siRNA are applied to the cells. After electroporation with a single pulse, cells were growth in RPMI complete medium , then analyzed 48h later.

#### XTT assay

AML cell lines (50 × $10^3$ cells/100µl) were incubated in a 96-well plate with indicated concentration of HA344 and #231 for 24h or 48h. 50 µl of XTT reagent (sodium 3'-[1-(phenylaminocarbonyl)-3,4-tetrazolium]-bis (4–methoxy–6–nitro) benzene sulfonic acid hydrate) was added to each well as described previously.(ref) The absorbance of the formazan product, reflecting cell viability, was measured at 490 nm. Each assay was performed in triplicate.

#### Cell Death

After stimulation with different drugs, cells were stained with 4',6-diamidino-2-phenylindole (DAPI) (Sigma, D9542, Saint Louis, MO, USA) 1µg/ml and then analyzed using the MACSQuant® cytometer (Miltenyi biotec, Bergisch Gladbach, Germany).

#### Western blot

AML cells were lysed at 4°C in lysis buffer. Lysates were centrifuged at 10 000g for 10 min at 4°C and supernatants were supplemented with concentrated SDS sample buffer. A total of 30 µg of protein were separated on 10 to 14% polyacrylamide gel and transferred onto polyvinylidene difluoride (PVDF) membrane (Immobilon-P, Millipore, Bedford, MA, USA). After blocking non-specific binding sites, the membranes were incubated with specific antibodies, washed three times and finally incubated with HRP-conjugated antibody for 1 h at room temperature. Immunoblots were revealed using the enhanced chemiluminescence detection kit (Amersham Biosciences, Uppsala, Sweden).

### Nano Differential Scanning Fluorimetry

NanoDSF experiments were performed using a NanoTemper Prometheus NT.48 instrument. 10 µM enzyme was incubated with 100 µM compound or DMSO for 2h at room temperature before the measurements. Standard capillaries were filled and heated from 25 to 95 °C with a ramp rate of 4.0 °C/min. Three technical replicates were carried out for each condition, and their means and standard deviations are depicted. The measured fluorescence ratio of the detected fluorescence at 350 nm and 330 nm was plotted as a function of temperature and the melting temperature was calculated as the inflection point of the resulting sigmoidal curve.

### Analysis of retrospective data

mRNA quantification of GPX4 from the Microarray Innovations in Leukaemia study (**MILE**) was analyzed via bloodspot website ("bloodspot.eu"). This analysis enables the quantification of GPX4 mRNA in normal cells, cells from MDS and AML patients, according to the mutations and chromosomal abnormalities found. Analysis of the impact of mRNA GPX4 levels on patient survival was carried out using the oncolnc portal based on the Cancer Genome Atlas Program cohort (**TCGA**). (http://www.oncolnc.org/, Anaya J. 2016. OncoLnc: linking TCGA survival data to mRNAs, miRNAs, and lncRNAs. PeerJ Computer Science 2: e67 https://doi.org/10.7717/peerj-cs.67 ). Comparison of GPX4 protein expression levels in 39 human cancer types was carried out using the DepMap platform ( https://depmap.org/portal/).

### Statistics

All data are presented as the mean ± SEM of at least three independent determinations. P-values were determined using the Prism V10 software (GraphPad, La Jolla, CA, USA). Unless stated otherwise in the figure legend, comparisons of the different groups were made with the one-way ANOVA test with Bonferroni correction. P-values of 0.05 (*), 0.01 (**), 0.001 (***), 0.0001(****) were considered statistically significant.
