## Supplementary material for "Dual targeting of GPX4 and TXNRD1 triggers eradication of AML cells through induction of apoptosis and ferroptosis": Chemical Synthesis

### General experimental information

All organic solvents were purchased from commercial sources and used as received. All chemicals were purchased from Aldrich, Merck or Alfa Aesar and used without further purification; thin layer chromatography (TLC) was performed on precoated Merck 60 GF254 silica gel plates and revealed by immersing ($KMnO_4$), and detection by means of UV light at 254 nm. $^{13}C$ NMR is proton decoupled. The coupling constants (*J*) are given in Hertz (Hz). The signals are reported as follows: chemical shift, multiplicity (s = singlet, d = doublet, t = triplet, q = quadruplet, m = multiplet, dd = doublet of doublets, ddd = doublet of doublets of doublets, dt = doublet of triplets, bs = broad singlet), coupling constant (*J*), integration and assignment. $^{1}H$ and $^{13}C$ NMR spectra were recorded on a Bruker Advance 400 MHz or 500 MHz spectrometer; chemical shifts are given in ppm and referenced to the solvent residual peak ($CDCl_3$). The purity of the compound was assessed by HPLC analysis; JASCO PU-2089 apparatus equipped with a UV detector, Ascentis Express Suppelco column (5 µm, RP-C18) 4.6 mm × 100 mm; step-wise gradient water/acetonitrile (1 ‰ formic acid): 5/95 during 2 min then 5/95 to 95/5 in 9 min, then a plateau at 95/5 for 5 min, then 95/5 to 5/95 during 1 min and then a plateau at 5/95 during 3 min; flow: 2 mL/min. High resolution mass spectra (HRMS) were recorded on a ThermoFisher Q Exactive (ESI-MS) at the resolution of 140 000 at m/z 200.

### Synthetic pathway to **VG40** and **VG60**

For the preparation of **compound II**, see M. G. van der Horst *et al.*, *J. Am. Chem. Soc.* **2010**, *132*, 5236.

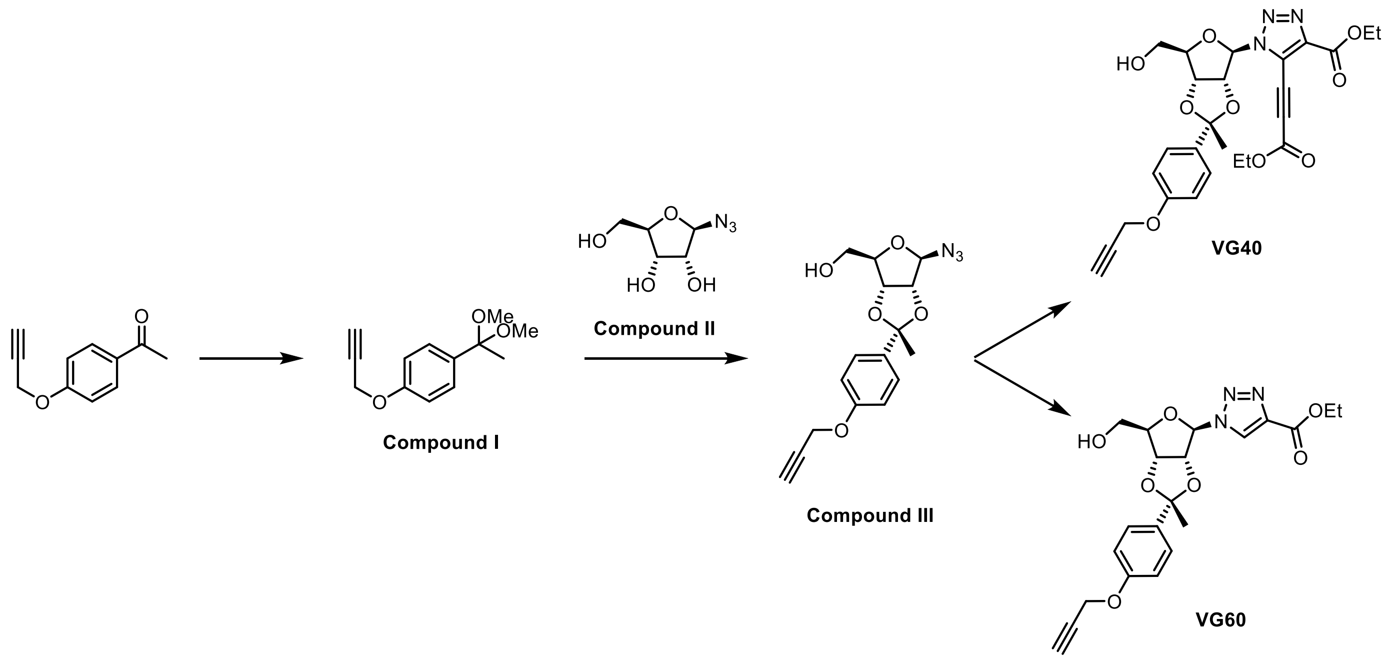


*Scheme S1: Synthesis of **VG40** and **VG60***

**Synthesis of Compound I**:

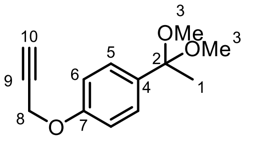


Commercially available 1-[4-(2-Propyn-1-yloxy)phenyl]ethanone (1.00 g, 5.75 mmol, 1.0 equiv.) and **p-toluenesulfonic acid** (100 mg, 0.58 mmol, 0.1 equiv.) were dissolved in anhydrous methanol (15 mL). Trimethylorthoformate (1.89 mL, 17.25 mmol, 3.0 equiv.) was added to the mixture and the reaction was allowed to stir at room temperature for 6 h. After complete consumption of the starting material (NMR monitoring in $CDCl_3$), the reaction mixture was quenched with triethylamine (0.16 mL, 1.15 mmol, 0.2 equiv.). Then, the solvent was evaporated under reduced pressure and the crude product (yellow oil, 1.485 g) was directly used for the synthesis of **compound III**; ***R*f** = 0.8 (Cyclohexane/EtOAc, 7/3); **$^1$H NMR (400 MHz, $CDCl_3$)** δ 7.42 (d, $^3J$ = 8.8 Hz, 2H, $H_5$), 6.95 (d, $^3J$ = 8.8 Hz, 2H, $H_6$), 4.69 (d, $^3J$ = 2.4 Hz, 2H, $H_{10}$), 3.17 (s, 6H, $H_3$), 2.52 (t, $^3J$ = 2.4 Hz, 1H, $H_{10}$), 1.52 (s, 3H, $H_1$).

**Synthesis of Compound III**:

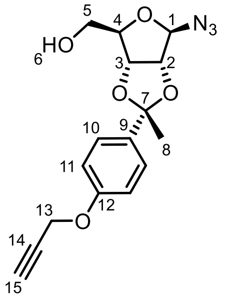


**Compound II** (502 mg, 2.87 mol, 1.0 equiv.) and **compound I** (1.265 g, 5.75 mmol, 2.0 equiv.) were dissolved in anhydrous *N,N*-dimethylformamide (2.6 mL). **p-toluenesulfonic acid** (24 mg, 0.14 mmol, 0.05 equiv.) was added to the mixture and the reaction was allowed to stir under vacuum for 16 h. Then, the reaction mixture was quenched with triethylamine (0.04 mL, 0.29 mmol, 0.1 equiv.) and diluted with EtOAc (50 mL). The organic layer was successively washed with 10 wt.% LiCl aqueous solution (50 mL) and brine (50 mL) before being dried over magnesium sulphate and concentrated to dryness in vacuo. The crude residue (brown oil, 2.105 g) was purified on a silica gel column chromatography using a mixture of Cyclohexane/EtOAc as the eluent (gradient: 10/0 to 8/2, v/v). Compound III was obtained as a yellow oil (227 mg, 24 %); ***R*f** = 0.3 (cyclohexane/EtOAc, 7/3); **$^1$H NMR (500 MHz, $CDCl_3$)** δ 7.45 (d, $^3J$ = 8.5 Hz, 2H, $H_{10}$), 6.95 (d, $^3J$ = 8.5 Hz, 2H, $H_{11}$), 5.47 (s, 1H, $H_1$), 4.91 (d, $^3J$ = 5.9 Hz, 1H, $H_2$), 4.70 – 4.68 (m, 2H, $H_{13}$), 4.66 (d, $^3J$ = 5.9 Hz, 1H, $H_3$), 4.34 (t, $^3J$ = 4.1 Hz, 1H, $H_4$), 3.72 (dd, $^2J$ = 12.2, $^3J$ = 3.3 Hz, 1H, $H_{5a}$), 3.66 (dd, $^2J$ = 12.0, $^3J$ = 4.6 Hz, 1H, $H_{5b}$), 2.53 (t, $^3J$ = 2.0 Hz, 1H, $H_{15}$), 2.30 (bs, 1H, $H_6$), 1.61 (s, 3H, $H_8$); **$^{13}$C NMR (126 MHz, $CDCl_3$)** δ 157.7 ($C_{12}$), 135.6 ($C_9$), 126.6 ($C_{10}$), 114.5 ($C_{11}$), 113.2 ($C_7$), 97.5 ($C_1$), 88.0 ($C_4$), 86.4 ($C_3$), 82.2 ($C_2$), 78.6 ($C_{14}$), 75.8 ($C_{15}$), 63.7 ($C_5$), 55.9 ($C_{13}$), 27.1 ($C_8$); **HRMS (ESI)** calcd. for $[M+H^+]$ $C_{26}H_{17}N_3O_6$ 332,1246; found: 332,1234.

**Synthesis of VG40;** ethyl 5-(3-ethoxy-3-oxoprop-1-yn-1-yl)-1-((2*R*,3a*R*,4*R*,6*R*,6a*R*)-6-(hydroxymethyl)-2-methyl-2-(4-(prop-2-yn-1-yloxy)phenyl)tetrahydrofuro[3,4-*d*][1,3]dioxol-4-yl)-1*H*-1,2,3-triazole-4-carboxylate:

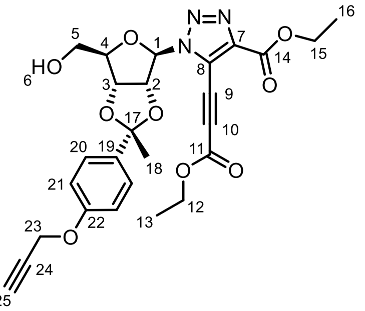

To a solution of **compound III** (75 mg, 0.23 mmol, 1.0 equiv.), in THF (2.8 mL) at 0°C, were successively added CuCN (24 mg, 0.27 mmol, 1.2 equiv.), ethylpropiolate (93 µL, 0.92 mmol, 4.0 equiv.), $H_2O_2$ (1.15 mmol, 100 µL of a 30% wt. solution in $H_2O$, 5.0 equiv.), and *N,N*-di*iso*propylethylamine (78 µL, 0.46 mmol, 2.0 equiv.). The reaction mixture was maintained at 0°C during 10 min. before being allowed to reach room temperature. Complete consumption of the starting azide was evidenced *via* thin layer chromatography (*ca.* 20 h, PE/EtOAc, 5/5 as the eluent). The reaction mixture was then filtered on calcite. The solid was washed with EtOAc and the filtrate was concentrated under vacuum. The crude material was purified over a silica gel column chromatography using a mixture of Cyclohexane/EtOAc as the eluent (gradient: 9/1 to 6/4 - v/v). VG40 was obtained as a brown oil (66 mg, 55%). $\boldsymbol{R_f}$ = 0.35 (PE/EtOAc, 5/5); **$^1$H NMR (500 MHz, CDCl$_3$)** δ 7.54 (d, $^3J$ = 8.4 Hz, 2H, $H_{20}$), 7.01 (d, $^3J$ = 8.3 Hz, 2H, $H_{21}$), 6.16 (d, $^3J$ = 2.4 Hz, 1H, $H_1$), 5.43 (dd, $^3J$ = 6.1, 2.4 Hz, 1H, $H_2$), 5.20 (dd, $^3J$ = 6.0, 2.0 Hz, 1H, $H_3$), 4.72 (d, $^3J$ = 2.2 Hz, 2H, $H_{23}$), 4.49 - 4.51 (m, 1H, $H_4$), 4.46 (q, $^3J$ = 7.1 Hz, 2H, $H_{15}$), 4.38 (q, $^3J$ = 7.3 Hz, 2H, $H_{12}$), 3.82 (dt, $^2J$ = 12.6, $^3J$ = 3.9 Hz, 1H, $H_{5a}$), 3.70 (ddd, $^2J$ = 12.8, $^3J$ = 8.5, 4.4 Hz, 1H, $H_{5b}$), 2.88 (dd, $^3J$ = 8.7, 4.9 Hz, 1H, $H_6$), 2.54 (t, $^3J$ = 2.4 Hz, 1H, $H_{25}$), 1.68 (s, 3H, $H_{18}$), 1.43 (t, $^3J$ = 7.1 Hz, 3H, $H_{16}$), 1.39 (t, $^3J$ = 7.4 Hz, 3H, $H_{13}$); **$^{13}$C NMR (126 MHz, CDCl$_3$)** δ 159.1 ($C_{14}$), 157.9 ($C_{12}$), 152.3 ($C_{11}$), 142.9 ($C_7$), 135.8 ($C_{19}$), 126.5 ($C_{20}$), 122.1 ($C_8$), 114.8 ($C_{21}$), 114.3 ($C_{17}$), 94.2 ($C_9$), 93.2 ($C_1$), 89.1 ($C_4$), 86.3 ($C_3$), 82.5 ($C_2$), 78.6 ($C_{24}$), 75.8 ($C_{25}$), 67.6 ($C_{10}$), 63.4 ($C_{12}$), 63.3 ($C_5$), 62.2 ($C_{15}$), 56.0 ($C_{23}$), 28.2 ($C_{18}$), 14.2 ($C_{13}$ or $C_{16}$), 14.2 ($C_{16}$ or $C_{13}$); **HPLC-UV ($\lambda_{254nm}$):** >99%; **HRMS (ESI)** calcd. for $[M+H^+]$ $C_{26}H_{28}O_9N_3$ 526,1826; found: 526,1820.

**Synthesis of VG60;** ethyl 1-((2*R*,3a*R*,4*R*,6*R*,6a*R*)-6-(hydroxymethyl)-2-methyl-2-(4-(prop-2-yn-1-yloxy)phenyl)tetrahydrofuro[3,4-*d*][1,3]dioxol-4-yl)-1*H*-1,2,3-triazole-4-carboxylate:

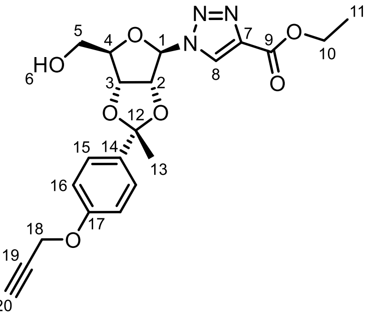

In a 4 mL vial, were successively charged **compound III** (86 mg, 0.26 mmol, 1.0 equiv.), $CuSO_4{\cdot}5H_2O$ (6 mg, 0.026 mmol, 10 mol%), ethyl propiolate (105 µL, 1.04 mmol, 4.0 equiv.) and a mixture of EtOH/$H_2O$ (1 mL, 1/1, v/v). Sodium ascorbate (10 mg, 0.052 mmol, 20 mol%) was then added and the reaction media was stirred at room temperature. Complete consumption of the starting azide was evidenced *via* thin layer chromatography (*ca.* 4 h., cyclohexane/EtOAc, 6/4 as the eluent). Next, the volatiles were evaporated in *vacuo* and the crude residue was purified over a silica gel column chromatography, using a mixture of Cyclohexane/EtOAc as the eluent (9/1 to 5/5, v/v). The title compound was obtained as a white oil (62 mg, 55%); ***R*****f** = 0.25 (cyclohexane/EtOAc, 6/4); **$^{1}$H NMR (500 MHz, $CDCl_3$)** δ 8.25 (s, 1H, $H_8$), 7.50 (d, $^{3}J$ = 8.8 Hz, 2H, $H_{15}$), 6.99 (d, $^{3}J$ = 8.8 Hz, 2H, $H_{16}$), 5.97 (d, $^{3}J$ = 2.2 Hz, $H_1$), 5.45 (dt, $^{3}J$ = 5.9, 2.3 Hz, 1H, $H_2$), 5.18 (dd, $^{3}J$ = 5.9, 1.6 Hz, 1H, $H_3$), 4.71 (d, $^{3}J$ = 2.4 Hz, 2H, $H_{18}$), 4.54 – 4.50 (m, 1H, $H_4$), 4.41 (q, $^{3}J$ = 7.1 Hz, 2H, $H_{10}$), 3.79 (dt, $^{2}J$ = 12.2, $^{3}J$ = 3.1 Hz, 1H, $H_{5a}$), 3.69 (ddd, $^{2}J$ = 11.9, $^{3}J$ = 6.8, 4.0 Hz, 1H, $H_{5b}$), 3.12 (dd, $^{3}J$ = 6.1 Hz, 4.5 Hz, 1H, $H_6$), 2.54 (t, $^{3}J$ = 2.4 Hz, 1H, $H_{20}$), 1.66 (s, 3H, $H_{13}$), 1.38 (t, $^{3}J$ = 7.1 Hz, 3H, $H_{11}$); **$^{13}$C NMR (126 MHz, $CDCl_3$)** δ 160.5 ($C_9$), 157.8 ($C_{17}$), 140.3 ($C_7$), 135.8 ($C_{14}$), 127.7 ($C_8$), 126.5 ($C_{15}$), 114.7 ($C_{16}$), 113.9 ($C_{12}$), 94.7 ($C_1$), 88.6 ($C_4$), 86.2 ($C_3$), 82.5 ($C_2$), 78.5 ($C_{19}$), 75.9 ($C_{20}$), 63.3 ($C_{11}$), 61.6 ($C_5$), 56.0 ($C_{18}$), 27.8 ($C_{13}$), 14.4 ($C_{10}$); **HPLC-UV ($\lambda_{254nm}$) :** 96 %; **HRMS (ESI)** calcd. for [M+H$^+$] $C_{21}H_{23}N_3O_7$ 430,1614; found: 430.1600.

### Copies of NMR spectra and HPLC chromatograms

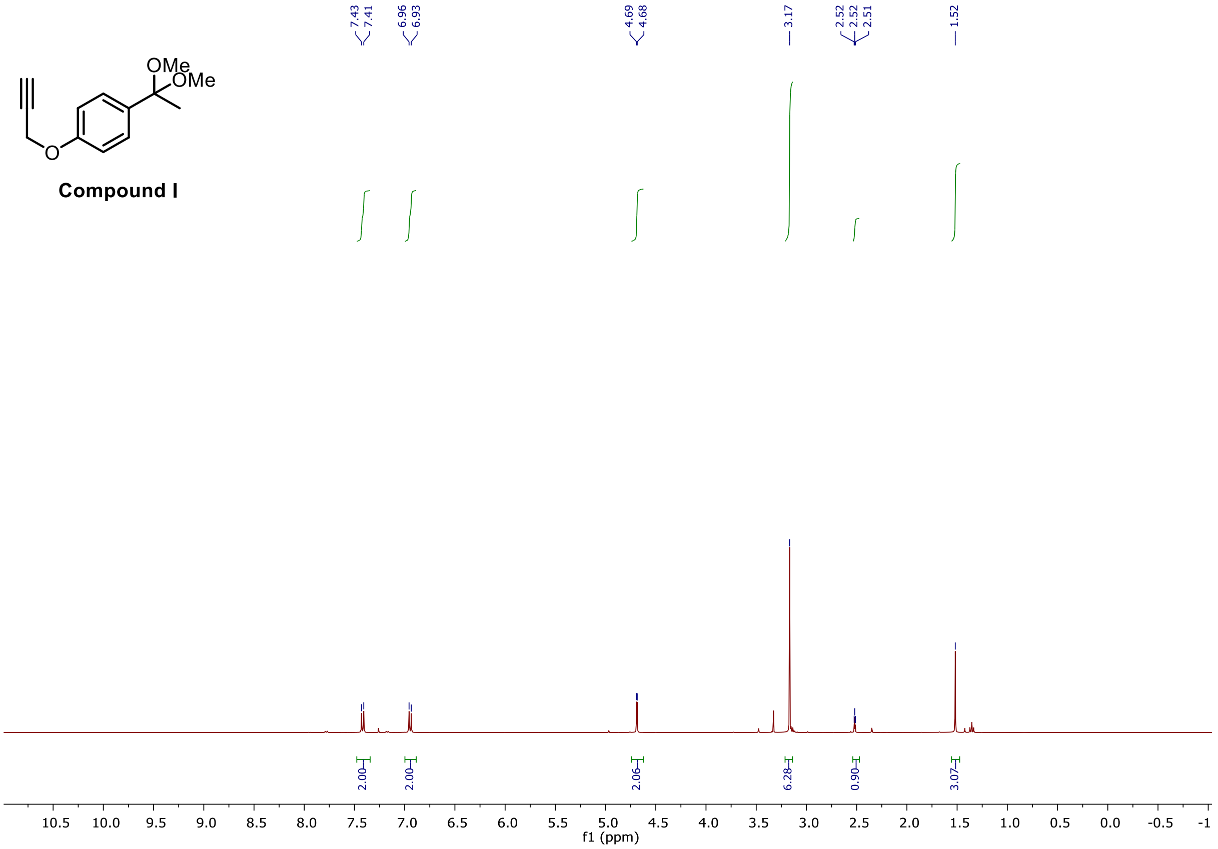


*Figure 1: $^{1}$H NMR (500 MHz, $CDCl_3$) spectrum of **compound I***

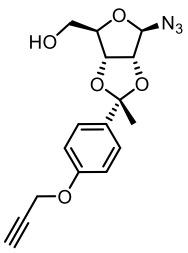

**Compound III**

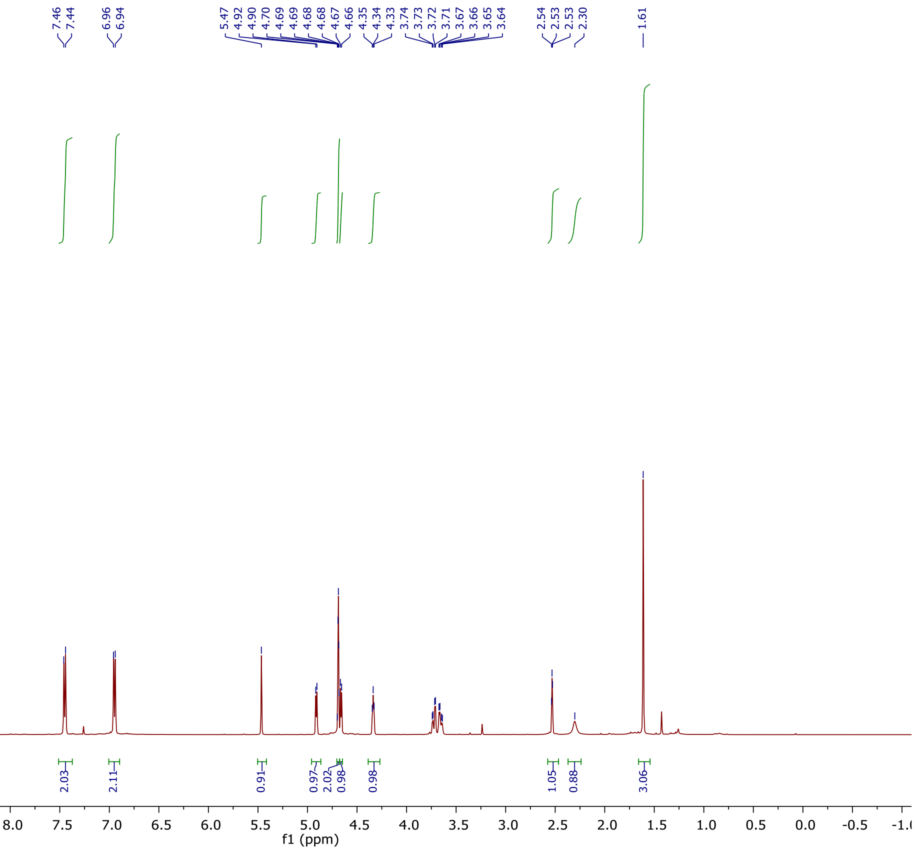


*Figure 2: ¹H NMR (500 MHz, $CDCl_3$) spectrum of **compound III***

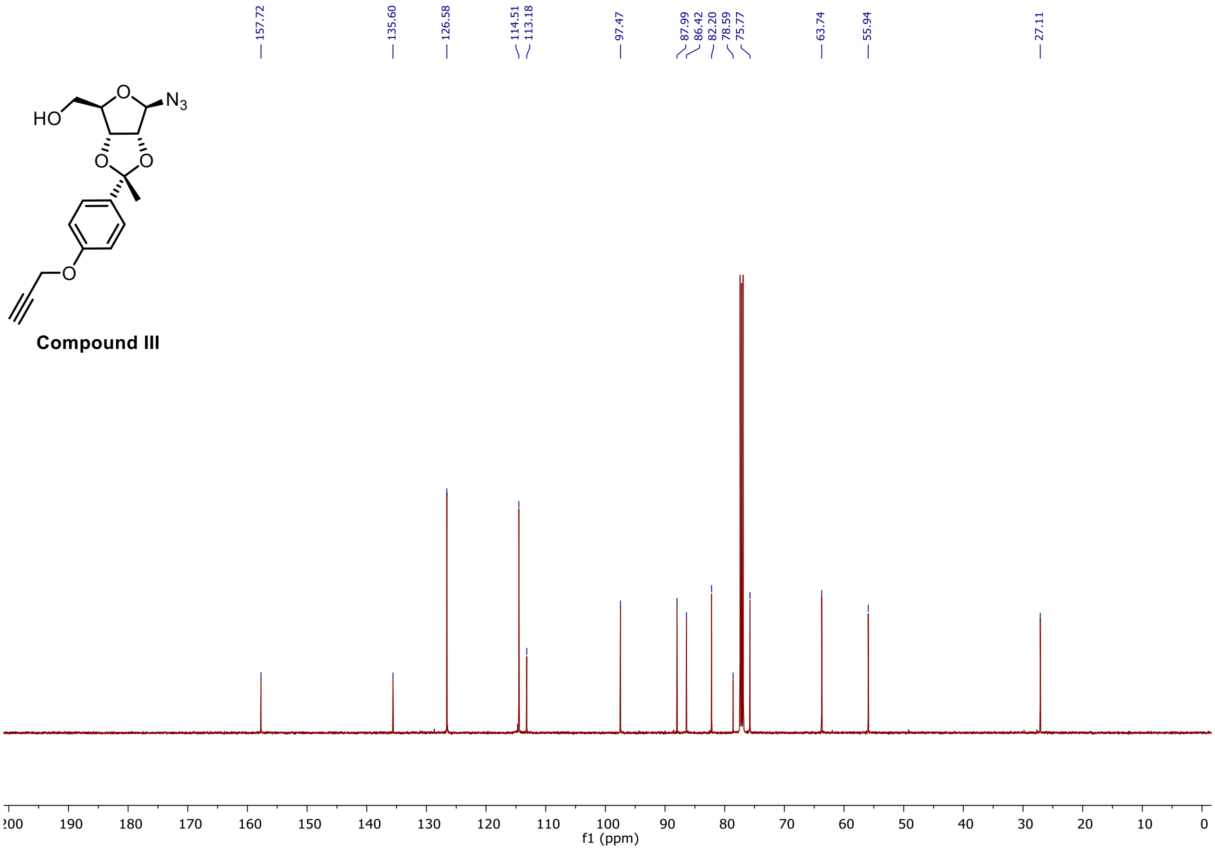


*Figure 3 : [13]C NMR (126 MHz, $CDCl_3$) spectrum of **compound III***

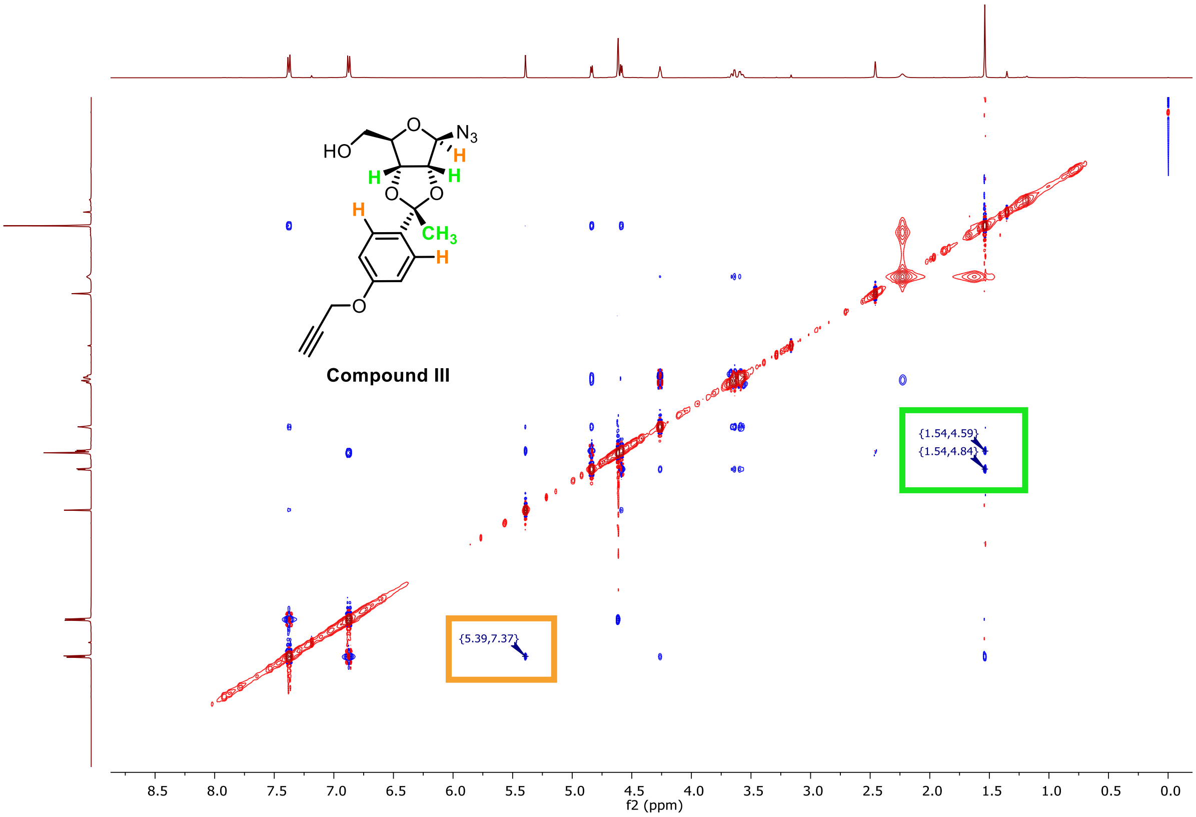


*Figure 4: [1]H NOESY NMR (500 MHz, $CDCl_3$) spectrum of **compound III***

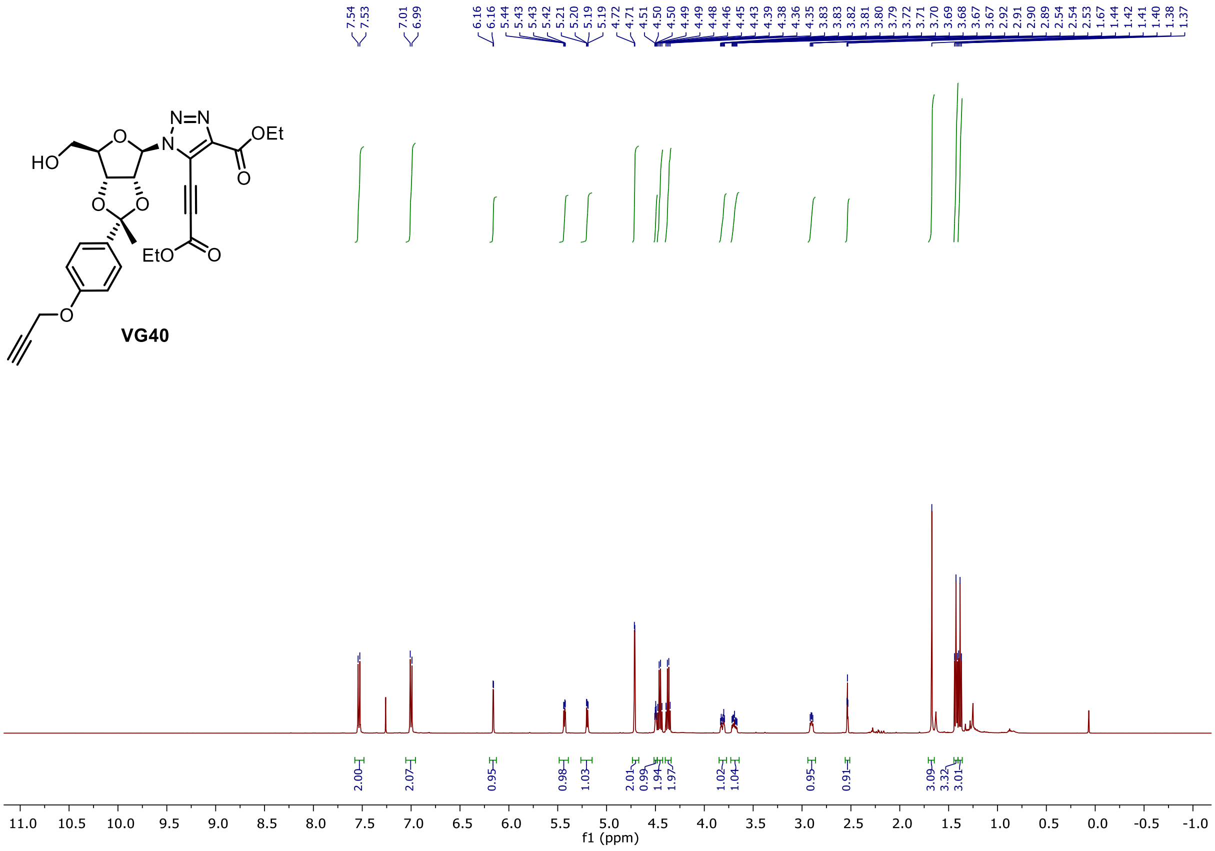


*Figure 5: $^{1}$H NMR (500 MHz, CDCl$_3$) spectrum of **VG40***

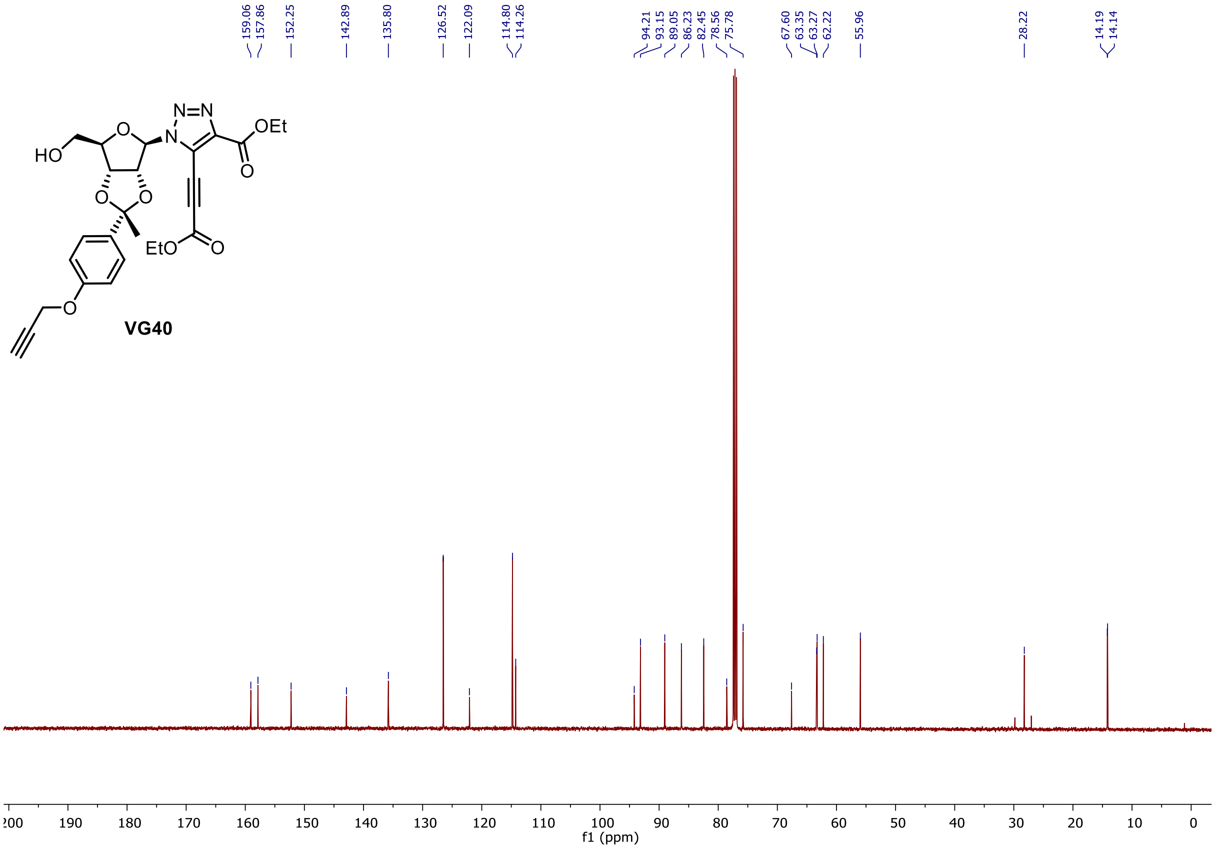


*Figure 6: $^{13}$C NMR (126 MHz, CDCl$_3$) spectrum of **VG40***

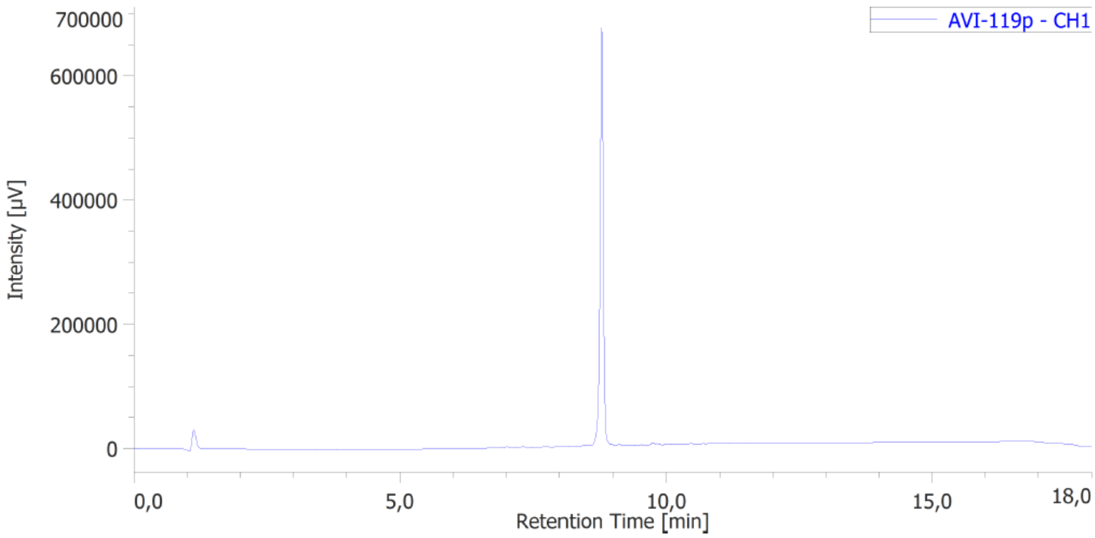


*Figure 7: HPLC chromatogram of **VG40***

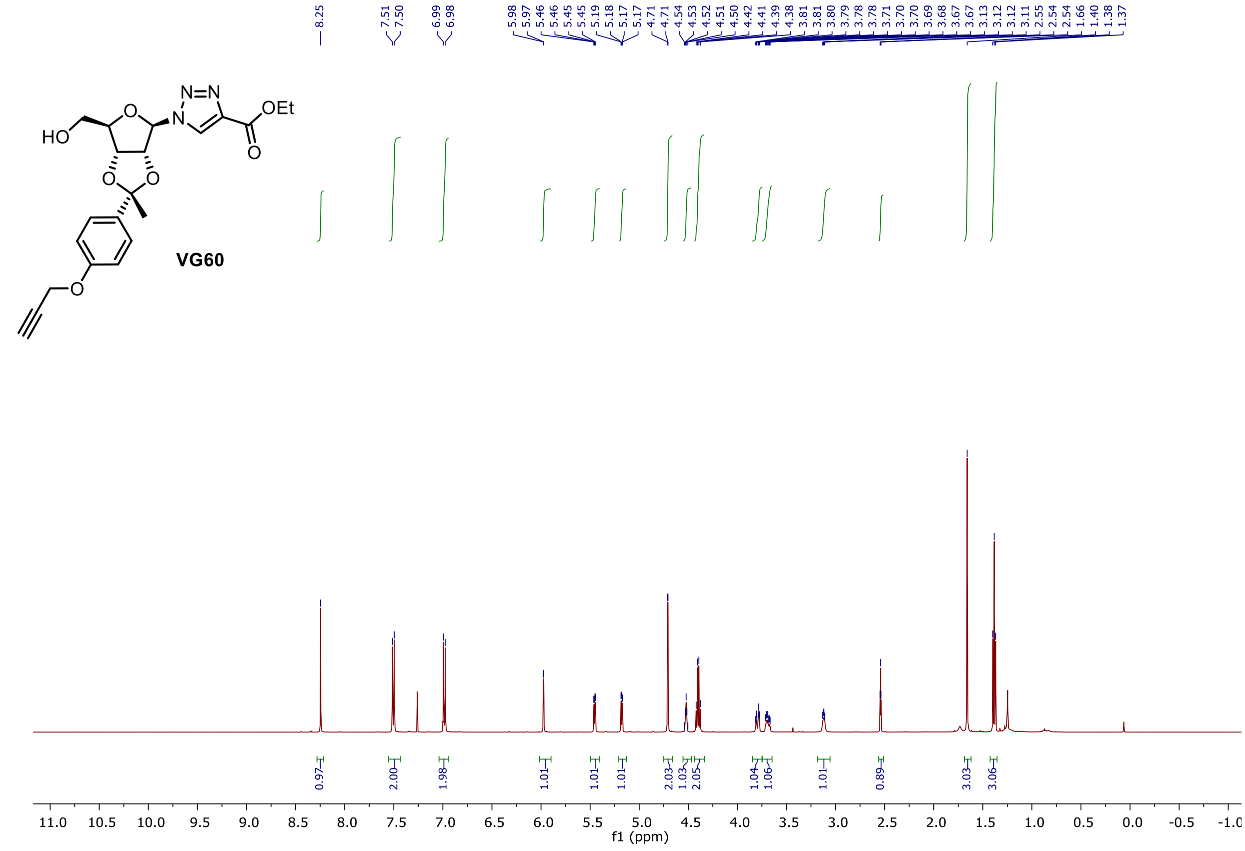


*Figure 8: [1]H NMR (500 MHz, CDCl$_3$) spectrum of **VG60***

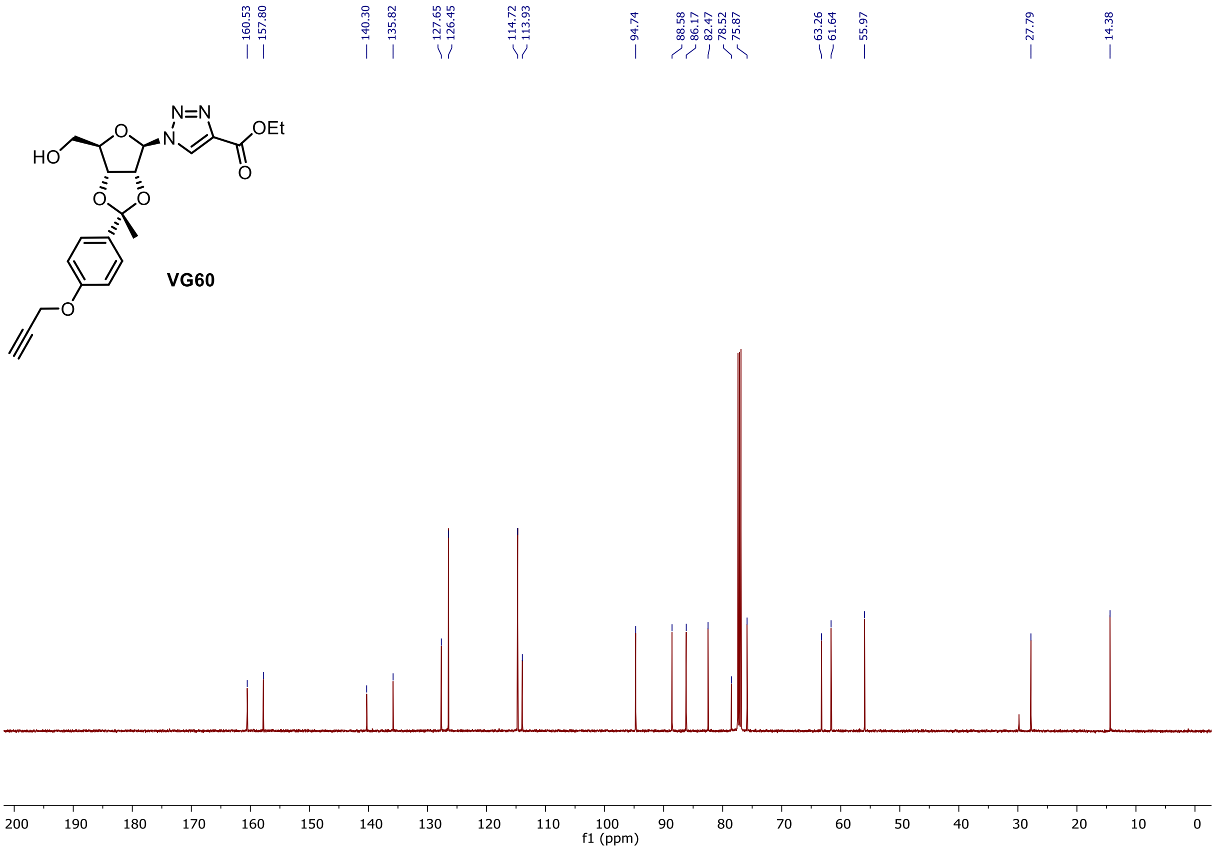


*Figure 9: $^{13}$C NMR (126 MHz, $CDCl_3$) spectrum of **VG60***

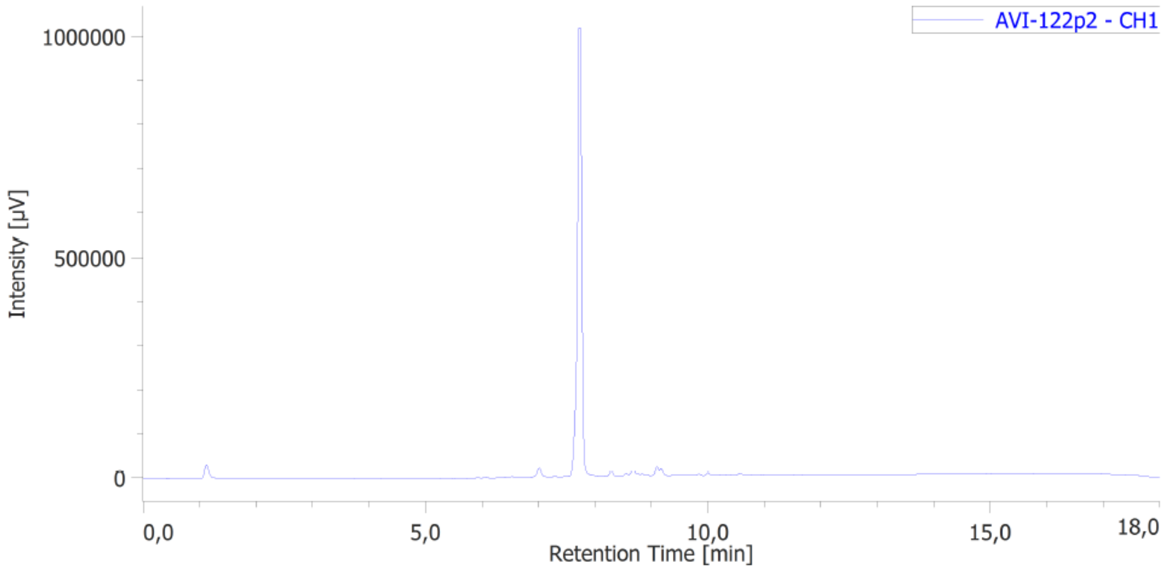


*Figure 10: HPLC chromatogram of **VG60***
